# A Multiscale Multicellular Spatiotemporal Model of Local Influenza Infection and Immune Response

**DOI:** 10.1101/2021.02.20.432089

**Authors:** T.J. Sego, Ericka D. Mochan, G. Bard Ermentrout, James A. Glazier

## Abstract

Respiratory viral infections pose a serious public health concern, from mild seasonal influenza to pandemics like those of SARS-CoV-2. Spatiotemporal dynamics of viral infection impact nearly all aspects of the progression of a viral infection, like the dependence of viral replication rates on the type of cell and pathogen, the strength of the immune response and localization of infection. Mathematical modeling is often used to describe respiratory viral infections and the immune response to them using ordinary differential equation (ODE) models. However, ODE models neglect spatially-resolved biophysical mechanisms like lesion shape and the details of viral transport, and so cannot model spatial effects of a viral infection and immune response. In this work, we develop a multiscale, multicellular spatiotemporal model of influenza infection and immune response by combining non-spatial ODE modeling and spatial, cell-based modeling. We employ cellularization, a recently developed method for generating spatial, cell-based, stochastic models from non-spatial ODE models, to generate much of our model from a calibrated ODE model that describes infection, death and recovery of susceptible cells and innate and adaptive responses during influenza infection, and develop models of cell migration and other mechanisms not explicitly described by the ODE model. We determine new model parameters to generate agreement between the spatial and original ODE models under certain conditions, where simulation replicas using our model serve as microconfigurations of the ODE model, and compare results between the models to investigate the nature of viral exposure and impact of heterogeneous infection on the time-evolution of the viral infection. We found that using spatially homogeneous initial exposure conditions consistently with those employed during calibration of the ODE model generates far less severe infection, and that local exposure to virus must be multiple orders of magnitude greater than a uniformly applied exposure to all available susceptible cells. This strongly suggests a prominent role of localization of exposure in influenza A infection. We propose that the particularities of the microenvironment to which a virus is introduced plays a dominant role in disease onset and progression, and that spatially resolved models like ours may be important to better understand and more reliably predict future health states based on susceptibility of potential lesion sites using spatially resolved patient data of the state of an infection. We can readily integrate the immune response components of our model into other modeling and simulation frameworks of viral infection dynamics that do detailed modeling of other mechanisms like viral internalization and intracellular viral replication dynamics, which are not explicitly represented in the ODE model. We can also combine our model with available experimental data and modeling of exposure scenarios and spatiotemporal aspects of mechanisms like mucociliary clearance that are only implicitly described by the ODE model, which would significantly improve the ability of our model to present spatially resolved predictions about the progression of influenza infection and immune response.

## Introduction

Respiratory viral infections continue to be a serious public health concern, from mild seasonal influenza strains to the highly pathogenic SARS-CoV-2 pandemic. In recent influenza strains associated with highly pathogenic outcomes, excess inflammation and cytokine storm tend to be major causes of mortality (1). Similarly, the recent COVID-19 epidemic has shown this coronavirus induces a similar cytokine storm in many of the lethal infections (2). A deeper understanding of the mechanisms involved in the initiation, proliferation, and reduction of the inflammatory response is key to understanding the reasons why some infections can become lethal.

Spatiotemporal dynamics impact nearly all components of the resolution of a viral infection both *in vitro* and *in vivo*. Viral replication rates, for example, depend on the type of cell the virus has invaded, the family and strain of the virus, and the strength of the immune response deployed against the pathogen. Viruses have been theorized to differ in replication rates between mucosal and bronchial epithelial cells (3). In addition, some viruses have been shown to localize to certain areas of the lungs rather than spread homogeneously throughout the respiratory tract. For instance, the 2009 pandemic H1N1 strain has been shown to replicate more extensively throughout the lower respiratory tract than either seasonal H1N1 or H5N1 (4). Seasonal H1N1 and H3N2 tend to replicate primarily in the bronchi, while H5N1 replicates largely in alveoli (5). In addition to these spatial differences, temporal differences in viral replication rates also play a part in differing levels of pathogenicity between viral strains. Multiple experimental studies have shown that strains exhibit distinct rates and mechanisms for cell entry, replication, and evasion of immune responses, allowing certain strains to be more virulent than others (5–8). The immune response to viral infection also includes many spatially-resolved biological processes, many of which are poorly understood, such as the search strategies of CD4^+^ and CD8^+^ T cells leading to antigen recognition, memory and effector T cell differentiation, migration via chemokinesis, chemotaxis and haptotaxis, and cytotoxic killing of infected cells (9). Heterogeneous spread of virus and infected cells has been theorized to affect the spread of infection through the lung; clusters of dead cells near productively-infected cells may prevent the virus from spreading (10). This effect has been seen after lethal H5N1 infection in ferrets (5); excessive damage in the lower respiratory tract prevents the virus from spreading further through the lung and limits the peak of the viral load. Thus, characterizing the spatial spread of the virus through the lung is critical to understanding the intrahost immune response to the infection.

Mathematical modeling has long been used to explore various details of the immune response to respiratory viral infections using ordinary differential equation (ODE) models (11– 14). However, spatial effects cannot be explored in a typical ODE model, as these models are founded on a well-mixed assumption that neglects spatially resolved biophysical mechanisms (*e*.*g*., lesion shape). Spatial models of the immune response have been developed in recent years to explore the effects of the spatial distribution of immune components on the resolution of infection (10,15–21). However, to our knowledge, no spatial model exists that describes host-pathogen interactions during influenza infection with cellular resolution while considering detailed descriptions of local and systemic aspects of both the innate and adaptive immune responses.

In this work, we combine the approaches of non-spatial ODE modeling and spatial, cell-based modeling to develop a multiscale, multicellular model of influenza infection and immune response. We generate much of our spatial model from a calibrated ODE model that describes infection, death and recovery of susceptible cells and innate and adaptive responses during influenza infection (14) using cellularization (22), a recently developed method for generating spatial, cell-based models from non-spatial ODE models. We develop models of cell migration and other mechanisms not explicitly described by the ODE model, and determine new model parameters to generate agreement between the spatial and original ODE models under certain conditions. We compare results between the models to investigate the nature of viral exposure and impact of heterogeneous infection on the time-evolution of the viral infection.

## Models and Methods

The ODE model of in-host response to influenza A virus from which the spatial model is generated describes infection and death of susceptible epithelial cells, and inflammatory, innate, adaptive and humoral responses. Population dynamics consist of explicit expressions for uninfected, infected and dead epithelial cells (*H, I* and *D*), macrophages (*M*), neutrophils in the blood and infected tissue (*Ñ* and *N*), antigen presenting cells (APCs, *P*), natural killer (NK) cells (*K*), B cells (*B*), and CD4^+^ and CD8^+^ T cells (*O* and *E*, respectively). Soluble signals of the model consist of tumor necrosis factor (TNF, *T*), interleukins 10 and 12 (IL-10 and IL-12, *L* and *W*, respectively), types I and II interferon (IFN, *F* and *G*, respectively), and generic chemokines (*C*), antibodies (*A*) and reactive oxygen species (ROS, *X*). We employ the method of cellularization (22) to generate a multiscale, multicellular, spatiotemporal model of local influenza A infection and immune response in an epithelial sheet. For details of the complete ODE model and cellularized spatial model, see *Appendix 1* in *Supplementary Materials*.

Cellularization describes the relationships of measurements of quantity at various scales of a biological system under well-mixed conditions. For a scalar quantity *Z* of a species at one scale, a scalar quantity Z of the same species at another scale, and a field distribution 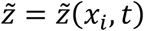 of which Z measures, according to cellularization,

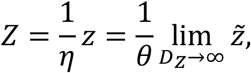

where *η* and *θ* are global and local scaling coefficients, respectively, and *D*_*Z*_ is the diffusion coefficient of 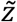 for diffusive species. For diffusive species, Z is the volume integral of 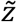 over a spatial domain, while for discrete objects of a particular type Z is the number of instances of the type of object (*e*.*g*., the number of neutrophils). Reaction-diffusion equations for locally heterogeneous soluble signals are generated from non-spatial descriptions. For a rate equation *Ż* = *v*(*Y, Z*) + *w*(*Y, Z*)*Q* for chemical species *Y* and *Z* and number *Q* of a cell type 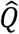, the analogous reaction-diffusion equation for 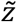 is

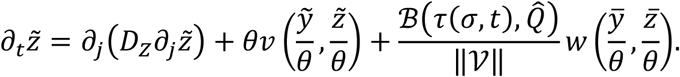

Here 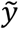 is the heterogeneous distribution associated with *Y*, 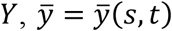 and 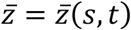 are the average value of 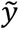 and 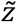 respectively, over the domain 𝒱 = 𝒱(*s, t*) of cell *s* with type τ(*s, t*), *σ* = *σ*(*x*_*i*_, *t*) = *s* at every site *x*_*i*_ occupied by cell *s* at time *t* (*i*.*e*., 𝒱(*s, t*) = {*x*_*i*_: *σ*(*x*_*i*_, *t*) = *s*}), and ℬ(*x, y*) is a Boolean-valued function equal to one when *x* = *y* and otherwise equal to zero.

Cellularization formulates cell-based stochastic dynamics using the Poisson cumulative distribution function from reaction kinetics that describe the inflow, outflow, and transitions by type (*e*.*g*., from alive to dead) of cell populations. For a number of cells *Q* of cell type 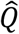 with mean inflow rate *f*, mean outflow rate *gQ*, and mean transition rate *uQ* to cell type *Ŝ* (*i*.*e*., 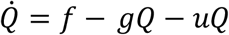 and 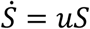 for *S* cells of type *Ŝ*) over a period [*t, t* + Δ*t*),

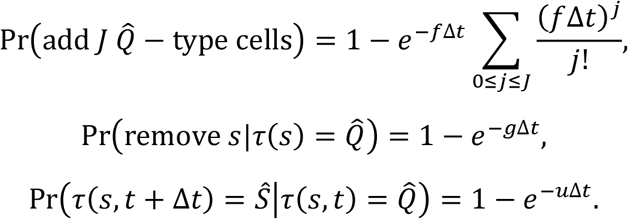

Local cell populations can be modeled such that a fraction of the population is explicitly modeled in a spatial domain, and the rest of the population act homogeneously. Cellularization describes the cell-based stochastic dynamics of a contact-mediated process with mean rate *r* (*i*.*e*., 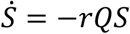 for *Q* and *S* cells of types 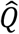 and *Ŝ* respectively) using an equipollent rate *γ* for cell *s* in an aggregate with total available contact surface area *A*_*U*_. If a process for a cell with total available contact surface area *A*_*s*_ is mediated by contact with a cell of type 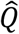 with total available contact surface area 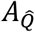, then for contact area 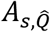 of cell *s* with a 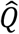 − type cell,

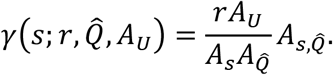

### Overview of Model Setup, Spatiotemporal Scaling and Cellularization

The milieu of the spatial, cell-based model is constructed by adapting an ODE model of influenza A infection and immune response (14) to comparable work on multiscale, spatial, cell-based modeling of viral infection and immune response (21) using cellularization. We consider a quasi-two-dimensional spatial domain in which local infection occurs in a fixed planar sheet of epithelial cells. Recruitment of various immune cell populations is governed by organismal-level dynamics coupled with signaling from within the spatial domain, where motile, locally acting immune cells are recruited from outside the spatial domain and placed on top of the epithelial sheet. Likewise organismal-level soluble signals are coupled with locally heterogeneous distributions of diffusive species in the spatial domain. We refer to model objects whose dynamics are explicitly modeled in the spatial domain as *local*, and likewise to those modeled with an ODE as *global*.

To model the spatial effects of infection, we model extracellular virus and uninfected, infected and dead cells as local (Figure 1). Type I IFN is modeled as local to model the spatial effects of antiviral resistance. Macrophages, chemokines and IL-10 are modeled as local to model the spatial effects of macrophage migration and diffusion in local inflammatory signaling. Likewise, NK and CD8^+^ T cells are modeled as local to model the effects of contact-mediated killing of infected cells in the immune response. B cells and blood neutrophils are not present at the local site of infection, and so are modeled as global. APCs primarily recruit other immune cell types according to the ODE model, and so we neglect the spatial aspects of their type I IFN release and model them as global. It follows that type II IFN, IL-12 and CD4^+^ T cells are modeled as global, since they immediately affect global objects. Antibodies, ROS and TNF could reasonably be modeled as local, however we assume that their rates of diffusion are sufficiently fast to approximate them as uniform throughout the simulation domain and model them as global. It follows that tissue neutrophils are modeled as global, since they release global ROS.

**Figure 1.**
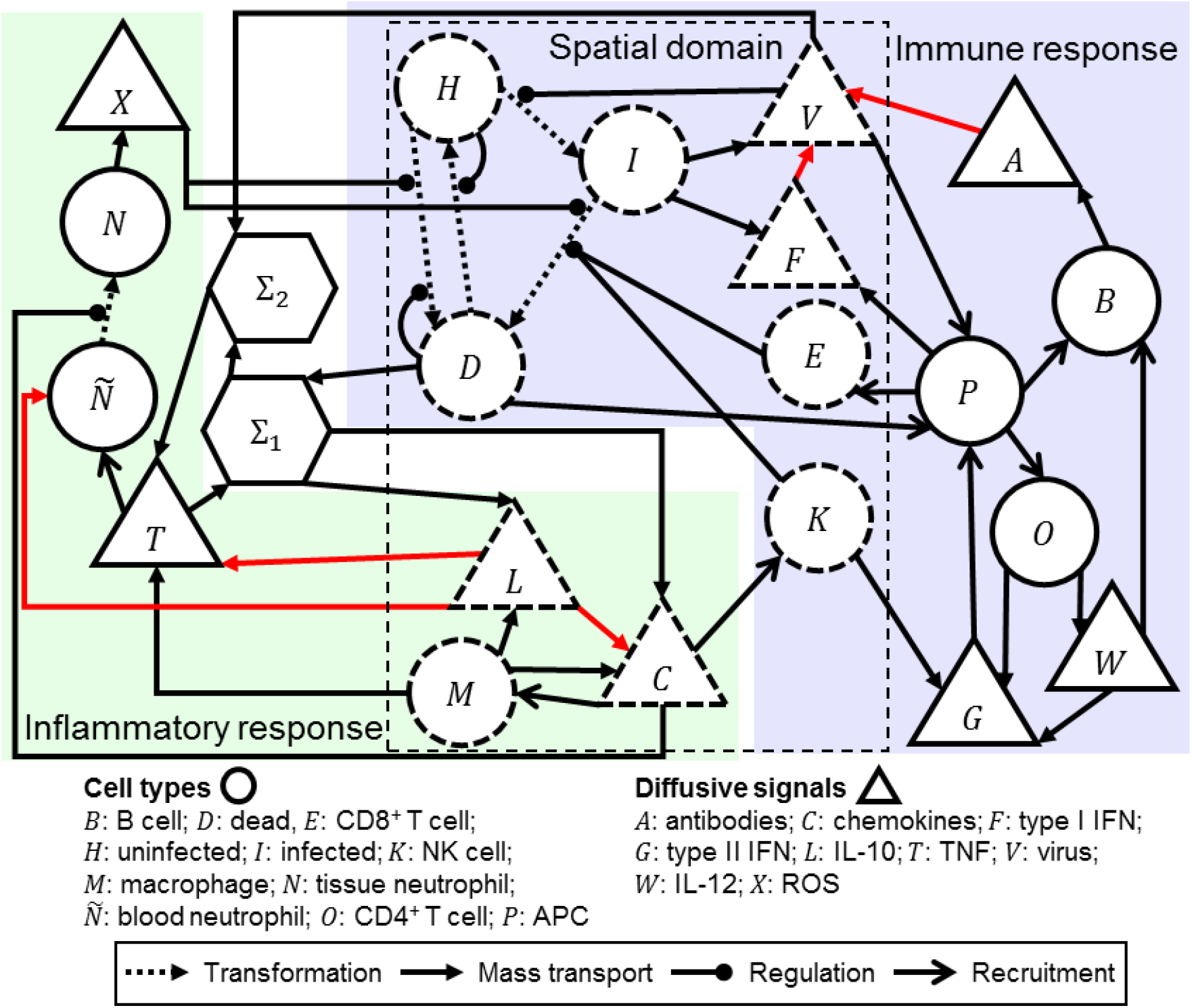
Schematic of the cellularizatized model of influenza infection and immune response. Model objects inside the dashed box labeled “Spatial domain” are modeled explicitly in the spatial domain, which are shown with dashed boundaries, whereas other model objects are treated as homogeneously acting due to their absence in the spatial domain (*e*.*g*., blood neutrophils) or their spatial properties (*e*.*g*., highly diffusive ROS) and are shown in solid borders. Analogous spatial and cell-based models of processes within, and across the boundary of, the spatial domain are derived from the ODE model using cellularization.

To model migration of local immune cells, we use the Cellular Potts model (CPM, or Glazier-Graner-Hogeweg model). The CPM is a lattice-based hybrid kinetic Monte Carlo method that represents generalized cells and medium as discrete, deformable, volume-excluding objects (23). Cell motility in the CPM is modeled as the stochastic exchanging of lattice sites by cells and medium according to minimization of a system effective energy ℋ, in this work written as,

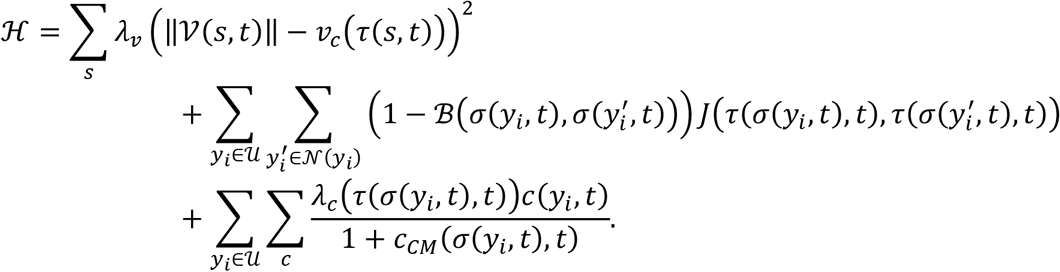

The first term implements a volume constraint *v*_*c*_ in each cell by cell type, the second term models adhesion at intercellular and cell-medium interfaces by cell type according to contact coefficients *J* using a neighborhood *𝒩*(*x*_*i*_) of each site, and the third term models logarithmic chemotaxis by cell type and field distribution according to a chemotaxis parameter λ_*c*_, local field concentration *c* and center-of-mass measurement *c*_*CM*_ of *c*. In this work we use a second-order Manhattan neighborhood for adhesion calculations, while applications of adhesion and chemotaxis modeling are described in the following section. In general, the CPM randomly selects a pair of neighboring lattice sites and considers whether the identification at one of the sites copies itself to the other site, called a copy attempt, which occurs with a probability according to a Boltzmann acceptance function,

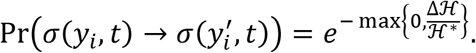

Here 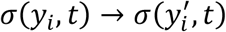 denotes the copy attempt where the identification at 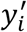 copies to *y*_*i*_, ℋ^∗^is the intrinsic random motility that affects the stochasticity of copy attempts, and Δℋ is the change in ℋ due to the copy. One simulation step, called a Monte Carlo step (MCS), consists of having considered a number of copy attempts equal to the total number of lattice sites.

### Particularities of the Cellularization

The ODE model defines a viral resistance *R* of the epithelial cell population due to the presence of type I IFN. Viral resistance affects a number of uninfected and infected cell behaviors, including decreased viral release and tissue recovery. Using cellularization, a cell-based viral resistance *ρ* = *ρ*(*s, t*) of each cell *s* with mean value of type I IFN in its domain 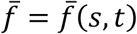 takes the form,

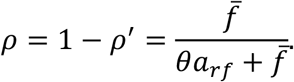

Here *a*_*rf*_ is a model parameter of the ODE model.

Diffusive transport is assumed to occur in a homogeneous medium, where for extracellular virus, chemokines, IL-10 and type I IFN we define the diffusion coefficients *D*_*V*_, *D*_*C*_, *D*_*L*_ and *D*_*F*_, respectively. The spatial model describes spatial and cell-based analogues of all mechanisms described by the ODE model for each heterogeneous species using partial differential equations (PDEs) of diffusive transport. Diffusive transport modeling of the extracellular virus 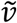 includes general decay, decay by the action of antibodies, mucociliary clearance, uptake by uninfected cells and release by infected cells; of chemokines 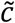 includes general decay, and release by macrophages regulated by TNF and the presence of dead cells; of IL-10 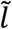 includes general decay, release by macrophages regulated by TNF and the presence of dead cells, and release by uninfected cells; of type I IFN 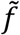 includes general decay, release by APCs, and release and uptake by infected cells (Table 1).

**Table 1.** PDEs generated from cellularization of the influenza ODE model for virus 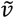, type I IFN 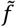, chemokines 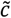 and IL-10 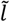 according to a general form for reaction-diffusion transport with diffusion coefficient. All symbols with subscripts are parameters from the ODE model. *η* and *θ* are the global and local scaling coefficients, respectively, according to cellularization. ℬ(*x, y*) is a binary function equal to one when *x* = *y* and zero otherwise. ‖𝒱(*σ, t*)‖ is the volume of a cell *σ* = *σ*(*x*_*i*_, *t*) at *x*_*i*_ and time *t* with type τ(*σ, t*) (*e*.*g*., uninfected *Ĥ*, infected *Î*, macrophage 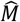). *a, d, p* and *T*^*^ are the total antibodies, dead cells, APCs and TNF in the spatial domain.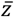 is a mean cellular measurement of 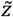

| General form | | $\partial_t \tilde{z} = (D_Z \partial_i^2 - q) \tilde{z} + r$ |
| --- | --- | --- |
| Field | Decay rate $q$ | Source rate $r$ |
| Virus $\tilde{v}$ | $\mu_v + \frac{g_{va}a}{\eta} + \frac{\theta g_v}{a_v \tilde{v} + \theta} + \frac{\mathcal{B}(\tau(\sigma, t), \hat{H}) g_{vh}}{\theta \ \mathcal{V}\ }$ | $\frac{\mathcal{B}(\tau(\sigma, t), \hat{I}) g_{vi} \rho'}{\ \mathcal{V}\ }$ |
| Type I IFN $\tilde{f}$ | $\mu_f + \frac{\mathcal{B}(\tau(\sigma, t), \hat{I}) g_{fi}}{\theta \ \mathcal{V}\ }$ | $\frac{\theta b_{fp} p}{\eta} + \frac{\mathcal{B}(\tau(\sigma, t), \hat{I}) b_{fi} \rho'}{\ \mathcal{V}\ }$ |
| Chemokines $\tilde{c}$ | $\mu_c$ | $\frac{\mathcal{B}(\tau(\sigma, t), \hat{M}) b_c}{\ \mathcal{V}\ \zeta}$ |
| IL-10 $\tilde{l}$ | $\mu_l$ | $\frac{1}{\ \mathcal{V}\ } \left( \frac{\mathcal{B}(\tau(\sigma, t), \hat{M}) b_l}{\zeta} + \mathcal{B}(\tau(\sigma, t), \hat{H}) \mu_l b_{lh} \rho' \right)$ |
| Auxiliary forms | $\zeta = 1 + \frac{\eta(g_1 \bar{l} + \theta g_2)}{(a_{11} T^* + a_{12} d)(\bar{l} + \theta d_2)}$ | $\bar{z} = \frac{1}{\ \mathcal{V}(\sigma, t)\ } \int_{\mathcal{V}(\sigma, t)} \tilde{z} dV$ |

The ODE model employs the Allee effect with a critical population of uninfected cells, above which recovery of uninfected cells occurs, and below which additional death of uninfected cells occur. We cellularize this mechanism by splitting it into individual stochastic events, of which a mean rate of death *a*_*D*_(*s*) due to the Allee effect is considered for each uninfected cell, and a mean rate of recovering a dead cell *a*_*H*_(*s*) is considered. The process of cell recovery is implemented as the transitioning of a dead cell to an uninfected cells (21). By treating both mechanisms of the Allee effect as contact-mediated and applying the well-mixed conditions (see *Appendix 1* in *Supplementary Materials* for derivations),

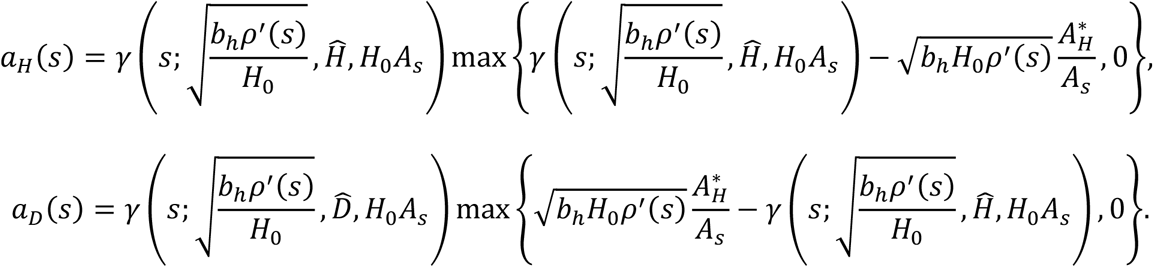

Here *b*_*h*_ is an ODE model parameter, *H*_0_ is the total number of epithelial cells according to the ODE model parameters and 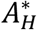 is a critical contact area with uninfected cells derived from the critical population value above which dead cells can recover, and below which uninfected cells can die. Using these forms and the cellularization of the remaining ODE model, the stochastic dynamics of the epithelial sheet occur according to the forms shown in Table 2, including infection of uninfected cells by extracellular virus, death of uninfected cells by ROS and the Allee effect, death of infected cells by ROS, contact-mediated killing by NK and CD8^+^ T cells, recovery of dead cells by the Allee effect and recruitment of local macrophages and NK and CD8^+^ T cells. Each type transition is considered once per simulation step for each cell in simulation, in the order of dead cells, infected cells, uninfected cells.

**Table 2.**
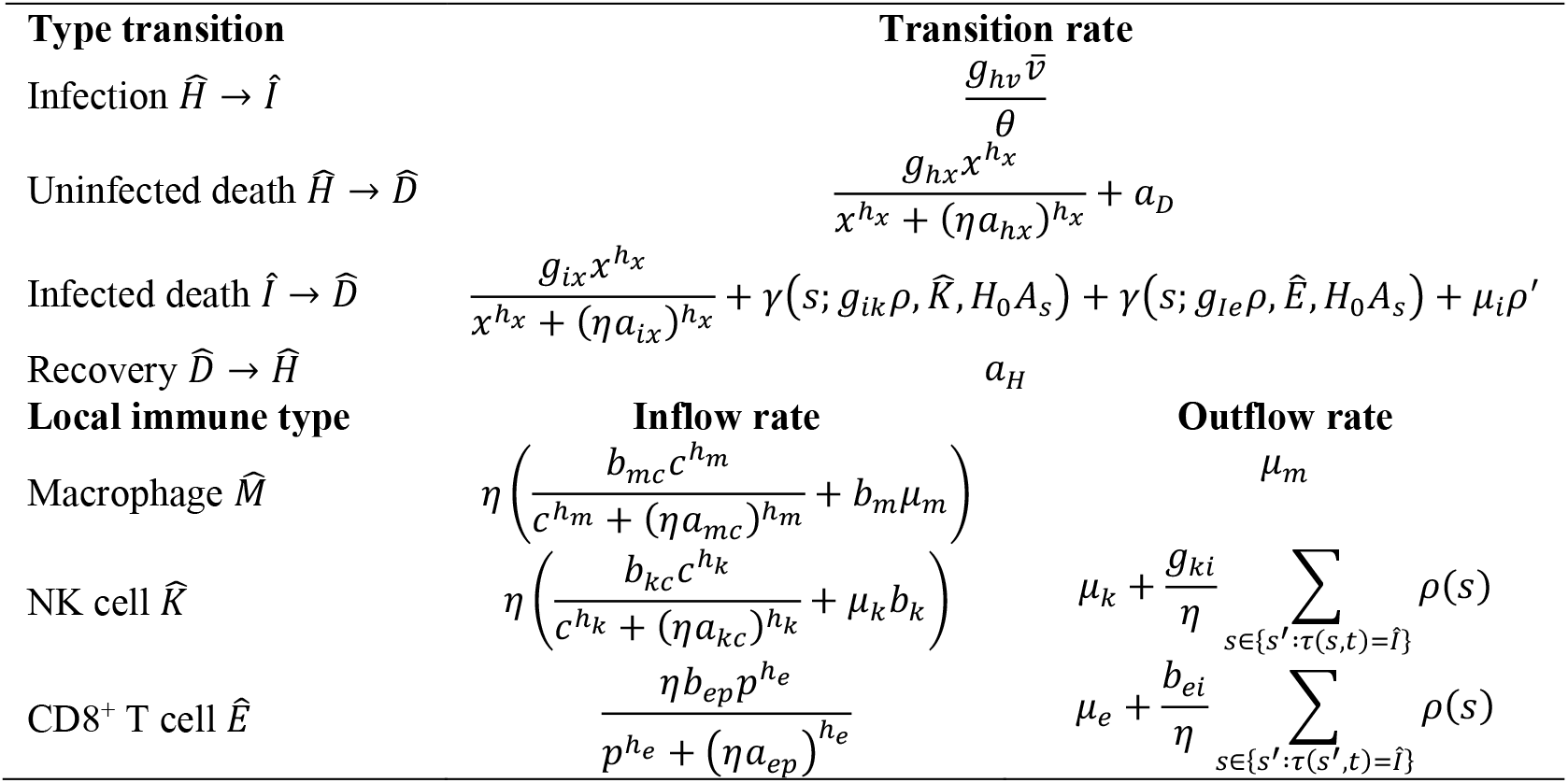
Stochastic dynamics of the epithelial sheet generated from cellularization of the influenza ODE model for epithelial cells of uninfected *Ĥ*, infected *Î* and dead 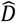 types and immune cells of macrophage 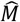, NK cell 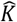 and CD8^+^ T cell *Ê* types. The transition from type *Ŷ* to type 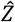 is denoted 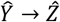. All symbols with subscripts are parameters from the ODE model. Mean cellular measurement of extracellular virus 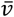 is calculated according to the form described in Table 1. *c, p* and *x* are the total chemokines, APCs and ROS in the spatial domain, and *ρ*(*s*) = 1 − *ρ*^′^(*s*) is the resistance of cell *s*.

| Type transition | Transition rate |  |
| --- | --- | --- |
| Infection $\hat{H} \rightarrow \hat{I}$ | $\frac{g_{hv}\bar{v}}{\theta}$ | |
| Uninfected death $\hat{H} \rightarrow \hat{D}$ | $\frac{g_{hx}x^{h_x}}{x^{h_x} + (\eta a_{hx})^{h_x}} + a_D$ | |
| Infected death $\hat{I} \rightarrow \hat{D}$ | $\frac{g_{ix}x^{h_x}}{x^{h_x} + (\eta a_{ix})^{h_x}} + \gamma(s; g_{ik}\rho, \hat{K}, H_0 A_s) + \gamma(s; g_{ie}\rho, \hat{E}, H_0 A_s) + \mu_i \rho'$ | |
| Recovery $\hat{D} \rightarrow \hat{H}$ | $a_H$ | |
| **Local immune type** | **Inflow rate** | **Outflow rate** |
| Macrophage $\hat{M}$ | $\eta \left( \frac{b_{mc}c^{h_m}}{c^{h_m} + (\eta a_{mc})^{h_m}} + b_m \mu_m \right)$ | $\mu_m$ |
| NK cell $\hat{K}$ | $\eta \left( \frac{b_{kc}c^{h_k}}{c^{h_k} + (\eta a_{kc})^{h_k}} + \mu_k b_k \right)$ | $\mu_k + \frac{g_{ki}}{\eta} \sum_{s \in \{s': \tau(s,t)=\hat{I}\}} \rho(s)$ |
| CD8<sup>+</sup> T cell $\hat{E}$ | $\frac{\eta b_{ep}p^{h_e}}{p^{h_e} + (\eta a_{ep})^{h_e}}$ | $\mu_e + \frac{b_{ei}}{\eta} \sum_{s \in \{s': \tau(s',t)=\hat{I}\}} \rho(s)$ |

### Additional Spatial Mechanisms

Beyond the cell-based models that can be generated from the ODE model using cellularization, the ODE model implicitly describes spatiotemporal aspects of influenza A infection and immune response that we can infer, impose or propose using additional data, assumptions and hypotheses. For the simplest case, employing the CPM requires imposing a volume constraint on each cell, the quantities, but not geometries, of which the ODE model describes. As such, we impose an approximate cell diameter of 10 µm on all cells according to the typical size of epithelial cells and simplification of negligible differences in typical volume among cell types (Table 3).

**Table 3.** Model parameters of spatial mechanisms used for all simulation.

| Parameter | Value | Source/Justification |
| --- | --- | --- |
| Volume constraint $v_c$ | 100 $\mu\text{m}^2$ | Chosen for an average cell diameter of 10 $\mu\text{m}$ |
| Volume multiplier $\lambda_v$ | 9 | (21) |
| *Diffusion coefficients* |  |  |
| Extracellular virus | 0.0119 $\mu\text{m}^2 \text{s}^{-1}$ | Chosen for a diffusion length of 5 cell diameters (21) |
| Chemokines | 1.04 $\mu\text{m}^2 \text{s}^{-1}$ | Chosen for a diffusion length of 10 cell diameters (21) |
| Type I IFN | 0.520 $\mu\text{m}^2 \text{s}^{-1}$ | Chosen for a diffusion length of 2 cell diameters for short-range anti-viral signaling |
| Interleukin-10 | 0.327 $\mu\text{m}^2 \text{s}^{-1}$ | Chosen for a diffusion length equal to that of chemokines |
| *Chemotaxis parameters $\lambda_c$* |  |  |
| Macrophage – virus | 5,000 | Chosen for strong chemotaxis according to typical field values |
| NK – chemokines | 5,000 | Chosen for moderate chemotaxis according to typical field values |
| CD8<sup>+</sup> T – chemokines | 10,000 | Chosen for strong chemotaxis according to typical field values |
| *Adhesion parameters $J$* |  |  |
| Uninfected – immune | 20 | Chosen for preferential attachment to infected cells |
| Infected – immune | 10 | Chosen for preferential attachment to infected cells |
| Dead – immune | 20 | Chosen for preferential attachment to infected cells |
| Homotypic immune | 25 | Chosen to prevent aggregation of immune cells |
| Heterotypic immune | 10 | Chosen to prevent aggregation of immune cells |

The ODE model describes the killing of infected cells proportionally to the number of NK and CD8^+^ T cells. In the spatial model, we place macrophages and NK and CD8^+^ T cells at the site of infection and explicitly model their shape and motility, which provides the opportunity to generate an explicit description of the spatiotemporal mechanisms involved in local immune cells locating and eliminating infection. We model immune cell locomotion by introducing chemotaxis and haptotaxis modeling to the biological objects and processes of the ODE model under the premise that NK and CD8^+^ T cells perform contact-mediated killing of infected cells, and that macrophages perform phagocytosis of virus and release soluble inflammatory signals (Figure 2). For a complete list of all behaviors, roles and properties of the cell types and fields of the model, see *Appendix 2* in *Supplementary Materials*.

**Figure 2.**
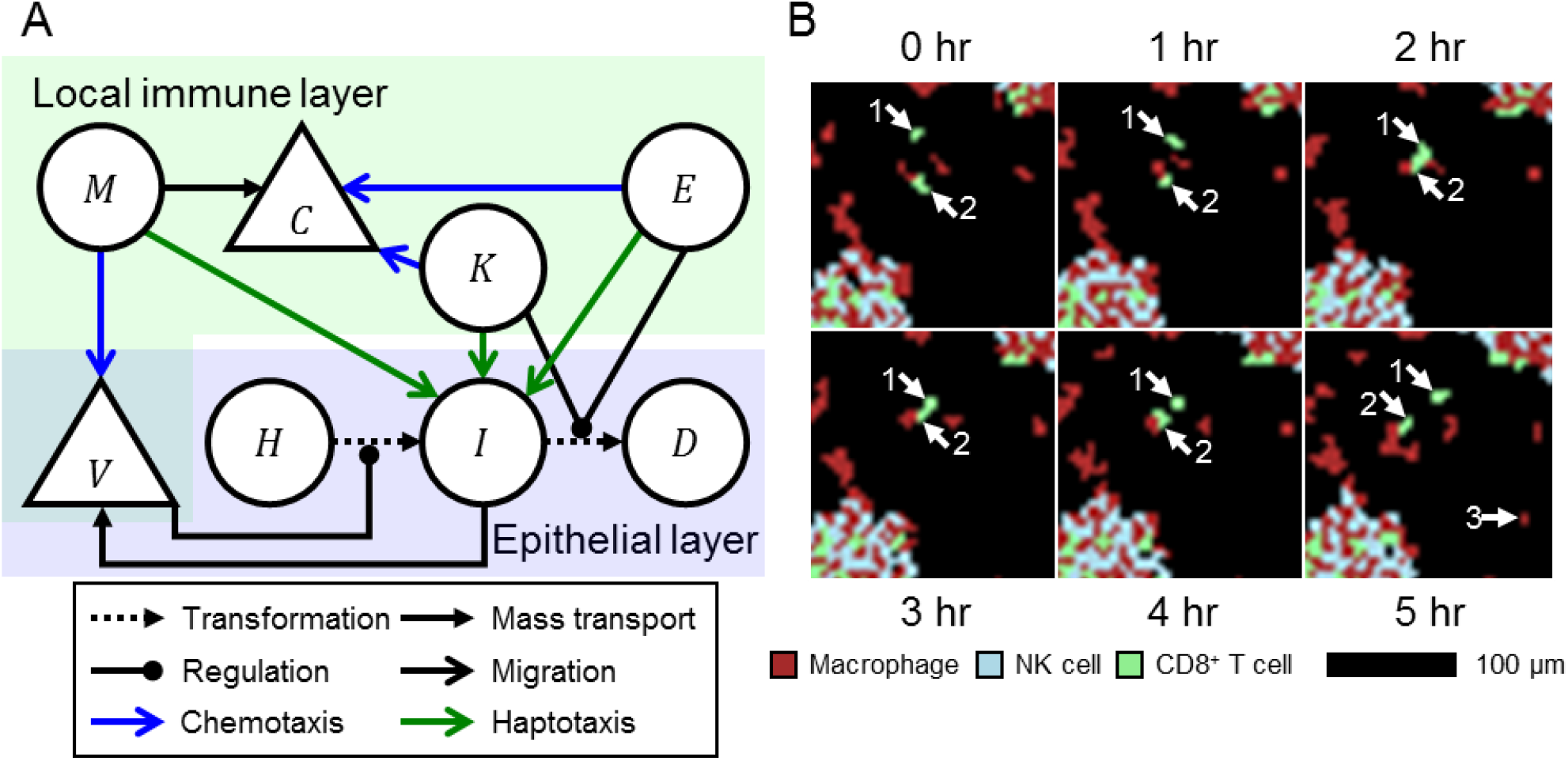
Local immune response model. A. Schematic of select model objects and processes in the spatial domain associated with infection, local immune response and local immune cell locomotion. Macrophages *M* chemotax towards extracellular virus *V* released by infected cells *I*, and release chemokines *C*. NK and CD8^+^ T cells *K* and *E* chemotax towards chemokines, and haptotax towards and kill infected cells through contact-mediated cytotoxic killing. B. Detailed view of local immune cells over six hours of simulation. Two CD8^+^ T cells (annotated “1” and “2”) migrate towards a common target, perform cytotoxic killing of underlying infected cells (not shown) and then migrate towards different targets. A macrophage is recruited to the local domain at hour 5 (annotated “3”). Aggregates of immune cells in the bottom left and top right of the detailed view respond to dense distributions of infected cells. Macrophages, NK cells and CD8^+^ T cells shown as maroon, cyan and green, respectively.

Based on phagocytosis and inflammatory signaling by macrophages, we model macrophages as chemotaxing up gradients of extracellular virus, and NK and CD8^+^ T cells as chemotaxing up gradients of chemokines. We also model the specialization of CD8^+^ T cells as their chemotactic sensitivity being twice that of NK cells. Based on contact-mediated killing of infected cells, we model stronger adhesion of immune cells to infected cells compared to uninfected and dead cells. We determined in early computational experiments that generating an effective local immune response also required preferential attachment that prevents excessive homotypic aggregation of immune cells but allows both heterotypic aggregates of immune cells and dispersion of immune cell aggregates according to chemoattractant distributions. As such, we model adhesion of immune cells to other immune cell types and the medium the same as to infected cells, and to immune cells of the same type the same as to uninfected and dead cells.

We approximated the diffusive characteristics of local soluble signals by diffusion length (*i*.*e*., √(*δ* /*q*) for diffusion coefficient δ and decay rate *q*) in units of cell diameters, using the decay parameters of the ODE model. Diffusion of extracellular virus and chemokines were approximated with diffusion lengths of five and ten cell diameters, respectively, based on previous, comparable modeling work on local infection and immune response (21). The diffusion length of IL-10 was assumed to be the same as that of chemokines, while type I IFN was modeled with a diffusion length of two cell diameters to model local anti-viral signaling.

### Implementation Details

Simulations were performed with comparable configurations to those in similar modeling work on local infection and immune response (21). All simulations were executed in CompuCell3D (24) with either a lattice planar dimension of 0.3 mm or 1.0 mm. Every lattice was discretized with a discretization length of 2 µm for cells that, on average, occupied 25 sites (Table 4). The local scaling coefficient *θ* = 4×10^−8^ µm^−2^ was calculated from the total number of epithelial cells according to the ODE model (250k) and cell volume constraint *v*_*c*_. The local scaling coefficient *η* was calculated as the ratio of the number of epithelial cells in the simulation domain to those in the ODE model parameters, and was 0.0049 and 0.04 for lattices with planar dimensions of 0.3 mm and 1.0 mm, respectively. Neumann and periodic conditions were applied to boundaries parallel and orthogonal, respectively, to the epithelial sheet. Epithelial cells were arranged in a uniform grid of 5×5 squares. All simulations used a time step of one minute per step, which was determined to be sufficiently small for numerically stability, particularly of type I IFN signaling. All ODE model parameters were taken from (14). Scaling was performed by epithelial cell population for an ODE model epithelial cell population of 250k cells. Using the prescribed volume constraint, simulations using only the ODE model showed the potential for local immune cells to exceed the available space in the immune cell layer, an event called overcrowding in cellularization. To mitigate overcrowding, we employed the cellularization strategy of partially homogenizing local immune cell populations, where all macrophages are explicitly represented in the spatial domain (since they provide directional signaling), while 25% of NK and CD8^+^ T cells act homogeneously as scalar-valued functions. All local immune cells were seeded into the immune cell layer with a seeding fraction of 1% according to field values of their chemoattactrants (*i*.*e*., by virus for macrophages, by chemokines for NK and CD8^+^ T cells).

**Table 4.** Implementation parameters used in all simulations.

| Parameter | Value | Source/Justification |
| --- | --- | --- |
| Time step $\Delta t$ | 1 min. step<sup>-1</sup> | Chosen for numerical stability |
| Lattice width | 2 $\mu\text{m}$ | Chosen for epithelial cell size of 5 x 5 sites |
| Intrinsic random motility $\mathcal{H}^*$ | 10 | (23) |
| Seeding fraction | 1% | Chosen for efficient local immune response (22) |
| *Local fraction* |  |  |
| Macrophages | 100% | Chosen to explicitly model directional signaling in immune response |
| NK cells | 75% | Chosen to mitigate overcrowding according to ODE model results |
| CD8<sup>+</sup> T cells | 75% | Chosen to mitigate overcrowding according to ODE model results |

## Results

In this section we present results from simulations of the spatial, cell-based model of influenza A infection and immune response. Given the stochasticity of the cell-based models, we simulate multiple simulation replicas for all initial conditions and parameter sets to demonstrate both their qualitative dynamical and stochastic features. In all scenarios, we employ one of two types of initial conditions: *initial viral load*, where simulations are initialized with a nonzero amount of virus, which is uniformly applied in the extracellular virus field; or *initial infection fraction*, where a fraction of epithelial cells are randomly selected at the beginning of simulation and initialized as infected. All replicas were executed for two weeks of simulation time at most, and were terminated early if all epithelial cells were dead (a determined lethal scenario), or if all were uninfected and total extracellular virus was less than 0.001, which was several orders of magnitude less than typical values during infection (an assumed non-lethal scenario). To compare results between the ODE and spatial models, we also simulated all scenarios using the ODE model while scaling results to the size of the spatial model.

### Testing Agreement in Small Epithelial Patches

To evaluate the agreement between the ODE and spatial models using the described cellularization in *Models and Methods*, we first simulated fifty replicas of small epithelial patches of area 0.3 mm x 0.3 mm with high initial infection fraction, which has been shown to mitigate potential spatial effects of initial infection in cellularized models of viral infection (22). As such, we imposed an initial infection fraction of 5% comparably to related previous work (25,26).

5% initial infection generated a lethal outcome in all simulation replicas within four days of simulation time, with marginal stochasticity among simulation replicas (Figure 3). Spatial distributions of local immune cells showed mostly sparsely distributed macrophages in the first day of simulation, with some aggregation near groups of infected cells. By one day of simulation time, NK and CD8^+^ T cells began arriving at the site of infection and accumulated in locations with high chemokines. By two days of simulation time, after most, if not all, epithelial cells had died, local immune cells formed branching patterns and intermixed by type. In general, simulation replicas well recapitulated ODE model results. Type I IFN results demonstrated the most notable differences around the time when the number of infected cells was at its maximum, with corresponding downstream effects. In particular, paracrine regulation of type I IFN release was inhibited by diffusion, which lead to greater production of type I IFN and, to a lesser extent, extracellular virus. Slightly greater extracellular virus resulted in slightly less antibodies and slightly earlier infection and death of all epithelial cells in some simulation replicas.

**Figure 3.**
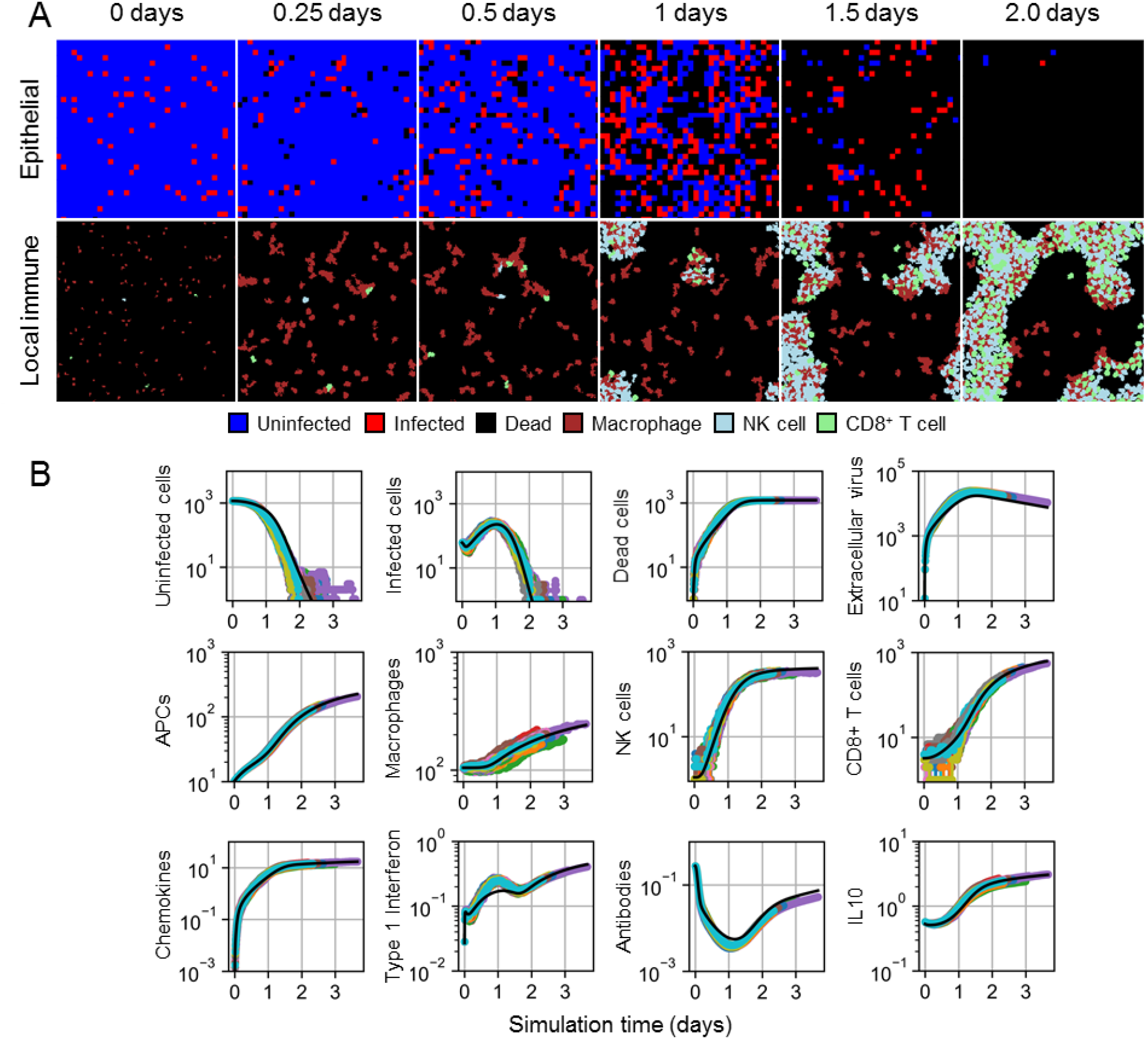
Spatial model results for 5% initial infection fraction. A. Spatial distribution of epithelial cells (top) and local immune cells (bottom) in a simulation replica at 0, 0.25, 0.5, 1, 1.5 and 2 days. Uninfected cells shown as blue, infected cells as red, dead cells as black, macrophages as maroon, NK cells as cyan, and CD8^+^ T cells as green. B. Results from 50 simulation replicas of the spatial model (colored lines) compared to ODE model results (black line) for epithelial cells, extracellular virus, and select immune cell types and signals.

### Disagreement in Large Epithelial Patches

Having shown acceptable agreement between the spatial and ODE models under marginally stochastic initial infection conditions, we generated a spatial equivalent of the lethal scenario to which the ODE model was calibrated by exposing large, uninfected epithelial patches to an initial viral load. We simulated 50 replicas of 1.0 mm x 1.0 mm epithelial patches, which, for the chosen spatial model parameters and 250k epithelial cells in the original ODE model simulations, collectively total two model organisms of the ODE model.

In all simulation replicas, at most, marginal infection occurred, in strong disagreement with ODE model predictions of a lethal outcome at around ten days (Figure 4). Figure 4A shows spatial distributions of one representative replica where any notable infection occurred, which consisted of an isolated lesion of infected and dead cells that recovered within two weeks. During early progression of such lesions, inflammatory signaling recruited significant numbers of macrophages, which localized at the lesion and subsequently recruited local NK and CD8^+^ T cells. Some new, later infection sites were also observed but well mitigated by present antibodies and quickly eliminated by the already stimulated immune response. In these cases, present local immune cells migrated with the general pattern of macrophages chemotaxing towards infected cells, followed by present, and reinforced by newly recruited, local NK and CD8^+^ T cells. In many other simulation replicas, no infection occurred, and the initial viral load decayed with no indication in the epithelial patch of exposure to virus (Figure 4B).

**Figure 4.**
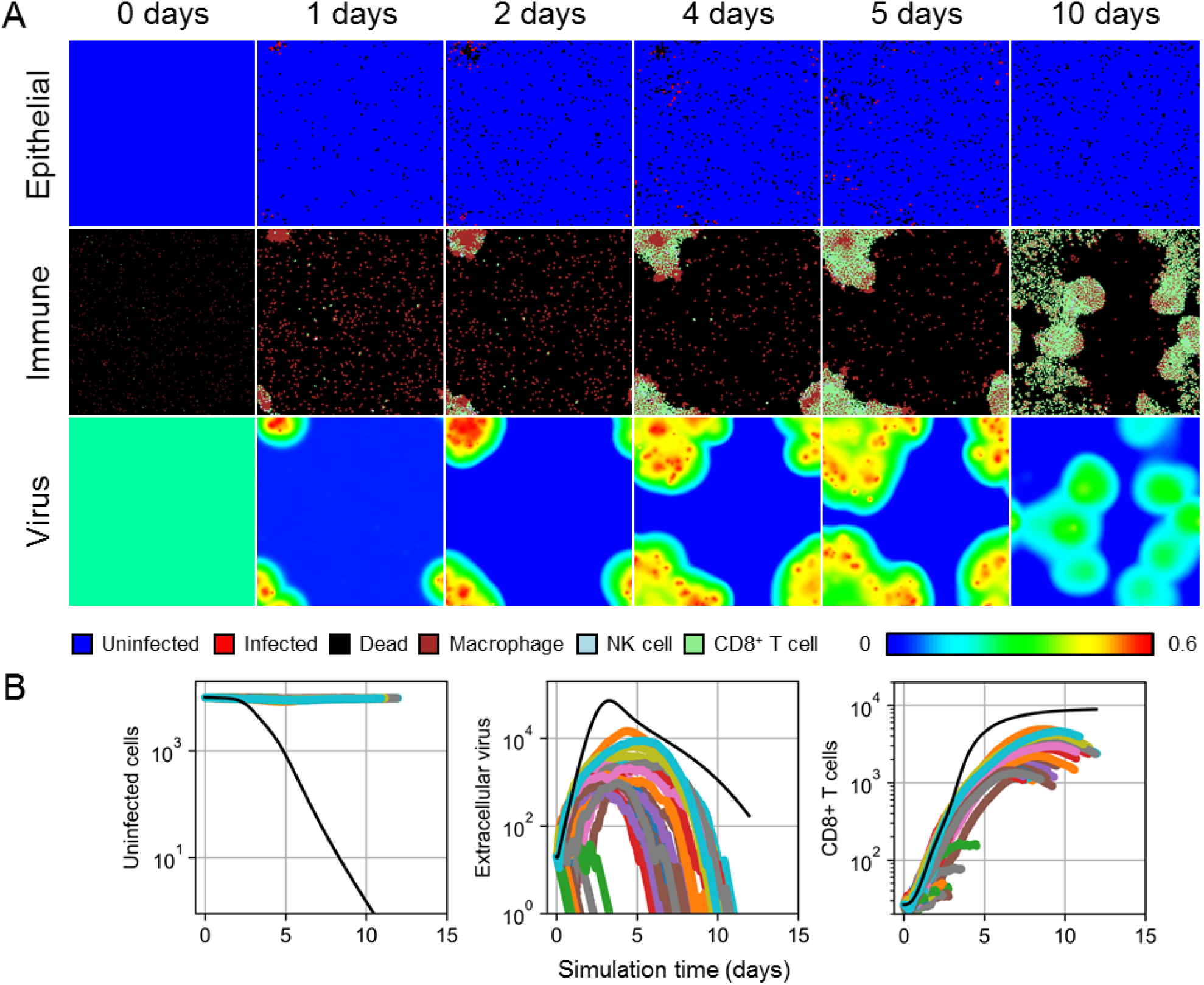
Spatial model results for the lethal exposure scenario. A. Spatial distribution of epithelial cells (top), local immune cells (middle) and extracellular virus (bottom) in a simulation replica at 0, 1, 2, 4, 5 and 10 days. Cell types shown as in Figure 3. The color bar shows contour levels of the extracellular virus distribution. B. Results from 50 simulation replicas of the spatial model (colored lines) compared to ODE model results (black line) for uninfected cells (left), extracellular virus (center), and CD8^+^ T cells (right).

### Only Large Initial Viral Load Produces Agreement

Because the initial viral load in the calibrated lethal scenario of the ODE model did not generate a lethal outcome in the spatial model, we tested varying initial viral loads to determine at what order of magnitude of initial viral load the spatial model generates a lethal outcome. Since the ODE model was calibrated to both non-lethal and lethal scenarios, where the lethal scenario differed from the non-lethal scenario only by a 10-fold increase in initial viral load, we performed a logarithmic parameter sweep of initial viral load by beginning with the spatial model equivalent to the non-lethal scenario, and increasing the initial viral load by a factor of 10 until the spatial model produced lethal outcomes in twenty simulation replicas.

We found that the spatial model begins to generate lethal outcomes when the initial viral load is at least greater than the initial viral load of the calibrated lethal scenario by a factor of 100 (Figure 5). Increasing the initial viral load of the lethal scenario (Figure 5, initial viral load of 10) by factors of 10 and 100 did not produce a lethal outcome in any simulation replica over two weeks of simulation time, though, besides the difference in outcome, the latter produced comparable predictions to those of the ODE model. A 1k increase in initial viral load from the lethal scenario produced at least nearly lethal outcomes in all simulation replicas, with the number of uninfected cells reaching minima at least as low as 10 cells (*i*.*e*., 0.1% uninfected). Many simulation replicas produced no uninfected cells at times as early as three and a half days, compared to about two and a half days according to the ODE model (*i*.*e*., when the number of uninfected cells is less than 1 according to the ODE model). In some replicas, marginal numbers of uninfected cells persisted as late as twelve and a half days, while two replicas demonstrated recovery of the epithelial patch and a corresponding non-lethal outcome. For this initial viral load, spatial model results disagreed otherwise only in amount of extracellular virus for replicas that produced a non-lethal outcome. We found these differences to be due to the difference in treatment of cell populations (*i*.*e*., as continuous quantities in the ODE model and as discrete quantities in the spatial model), where cell populations being less than one exhibits no notable effects in the ODE model, whereas discrete cell populations in the spatial model cease to exhibit any effects by having a number of cells equal to zero. For the calibrated non-lethal scenario (Figure 5, initial viral load of 1), no infection occurred in 19 out of 20 replicas, and in the one replica that did experience any infection, the maximum number of infected was two orders less, and occurred about two days earlier, than that of the ODE model.

**Figure 5.**
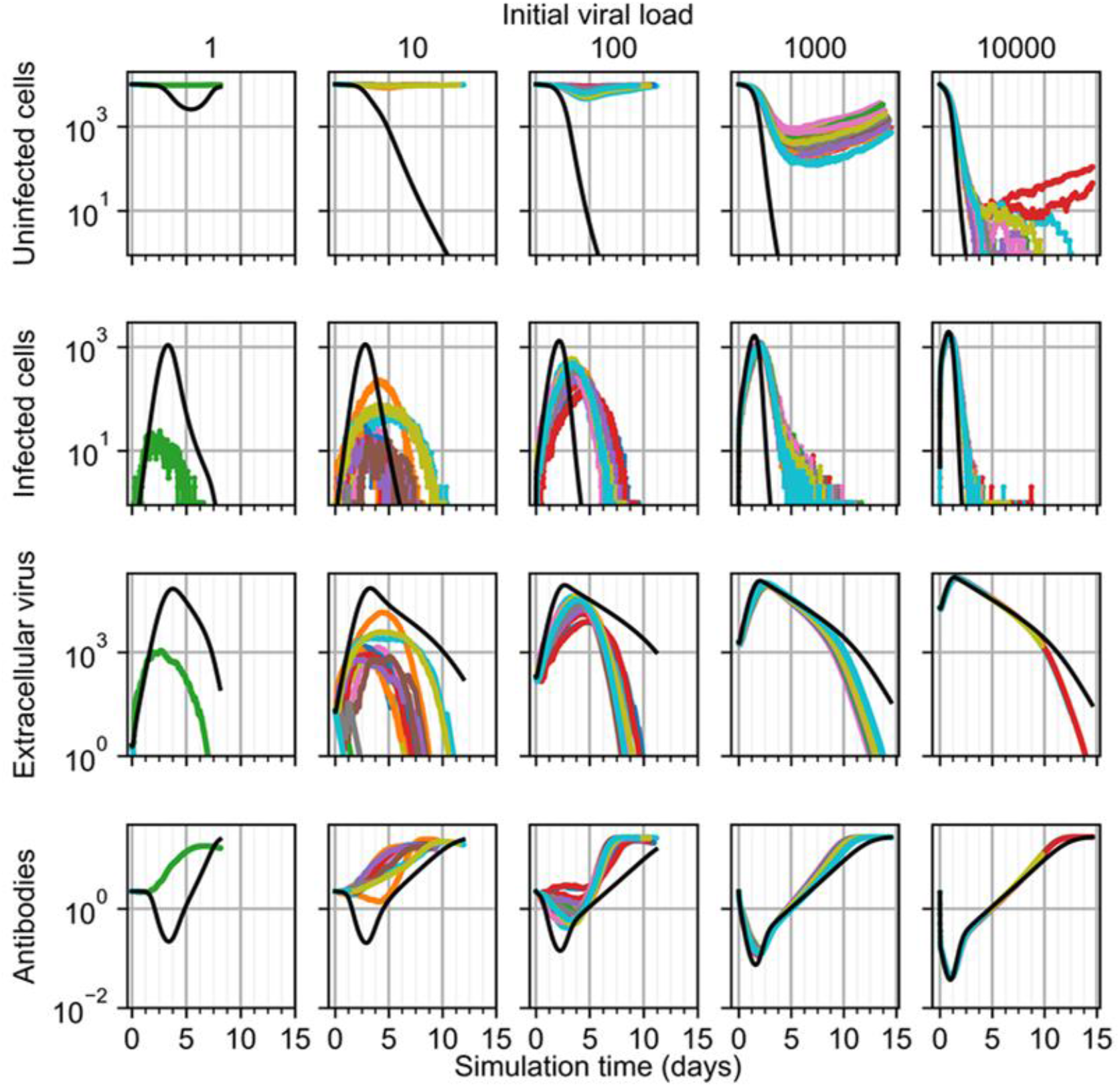
Results from simulation replicas of the spatial model (colored lines) compared to ODE model results (black line) for uninfected cells (top), infected cells (top-middle), extracellular virus (bottom-middle), and antibodies (bottom) for initial viral load multipliers, from left to right, of 1, 10, 100, 1000 and 10000.

For simulation replicas that did experience significant amounts of infection (*e*.*g*., for those with lethal outcomes), we observed multiple sites of significant infection within the first day after exposure (Figure 6). These sites were locations of significant recruitment of local macrophages and subsequent recruitment of local NK and CD8^+^ T cells, as well as localized type I IFN, which later became more homogeneous due to production by nonlocal APCs. Spatial distributions of chemokines and IL-10 showed gradients most apparently at around two days of simulation time, with IL-10 being greater in regions with significant accumulation of local immune cells, and became mostly homogeneous by around one week of simulation time when immune cells mostly covered the epithelial patch. For simulation replicas with the highest initial viral load that recovered (*e*.*g*., as in Figure 6), groups of uninfected epithelial that survived infection and immune response by around one week of simulation time became the sites of recovery of the epithelial patch, which became apparent by about two weeks of simulation time as outgrowths of uninfected cells into a distribution of otherwise dead cells.

**Figure 6.**
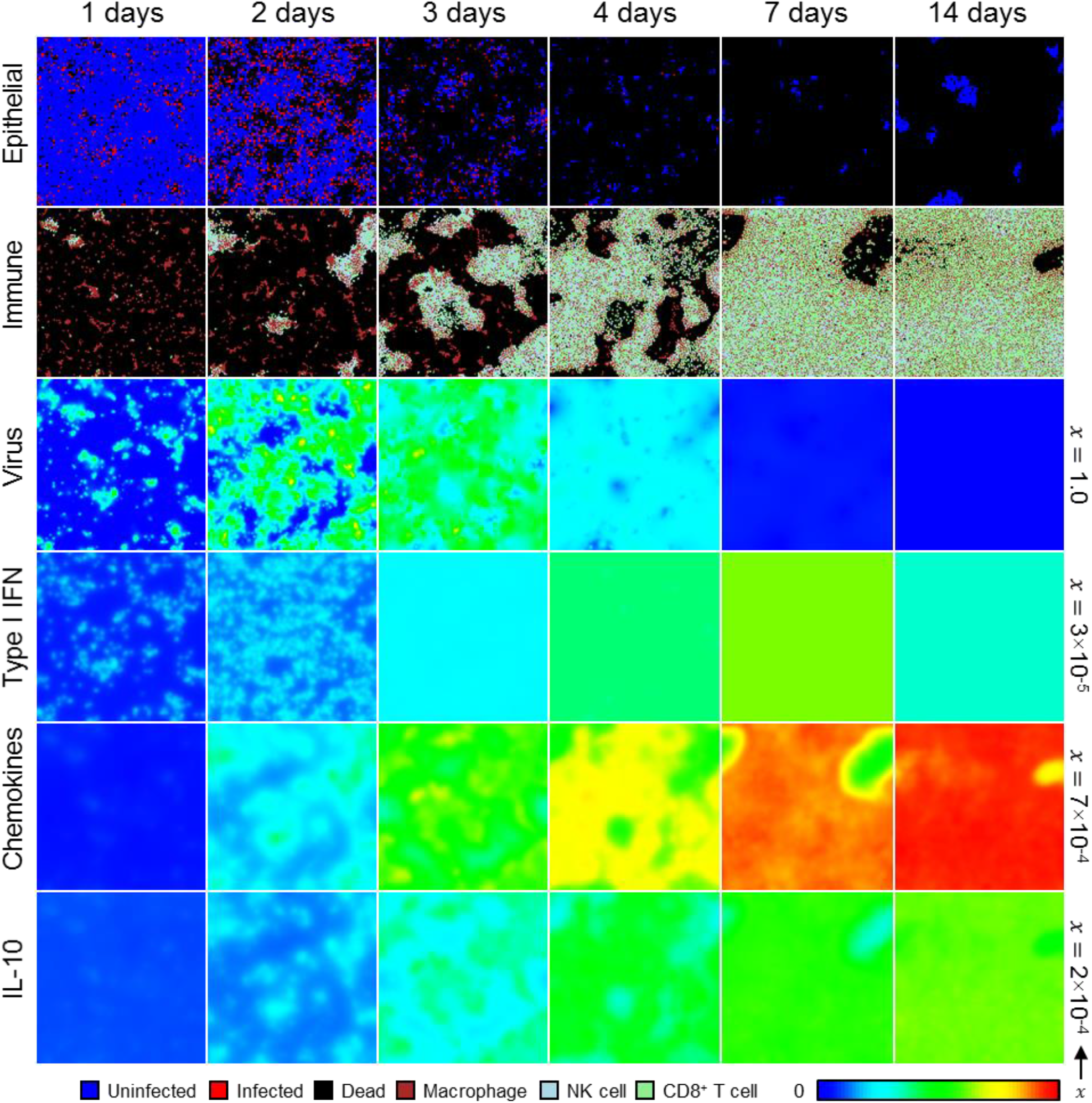
Spatial distribution of, from top to bottom, epithelial cells, local immune cells, and extracellular virus, type I IFN, chemokines and IL-10 in a simulation replica with 1000 initial viral load at 0, 1, 2, 4, 7 and 14 days. Cell types shown as in Figure 3. The color bar shows contour levels of diffusive species from zero to the maximum value per field. Each field maximum shown along the right border.

### Only Large Initial Fractions of Infected Cells Produce Agreement

Since much higher initial viral loads were required to generate significant infection in large epithelial sheets using the spatial model compared to the ODE model, we then tested agreement between the ODE and spatial models while varying initial infection fraction in 1 mm x 1 mm epithelial patches. We varied the initial infection fraction in a logarithmic sweep at intervals of 0.1%, 0.5%, 1% and 5% and simulated twenty simulation replicas for each initial infection fraction. As in *Testing Agreement in Small Epithelial Patches*, 5% initial infection fraction can produce fatal outcomes but in large epithelial sheets, though at times no earlier than about five days, but can also produce non-fatal outcomes (Figure 7). For all simulation replicas subjected to 5% initial infection fraction, the epithelial patch experienced infection comparably to that predicted by the ODE model, however in some cases a few uninfected cells survived and initiated recovery. As initial infection fraction decreased, peak extracellular virus and infected cells in the spatial model occurred later and with lesser magnitude, and all simulation replicas produced a non-fatal outcome for initial infection fraction less than or equal to 1%, all of which are fatal according to the ODE model.

**Figure 7.**
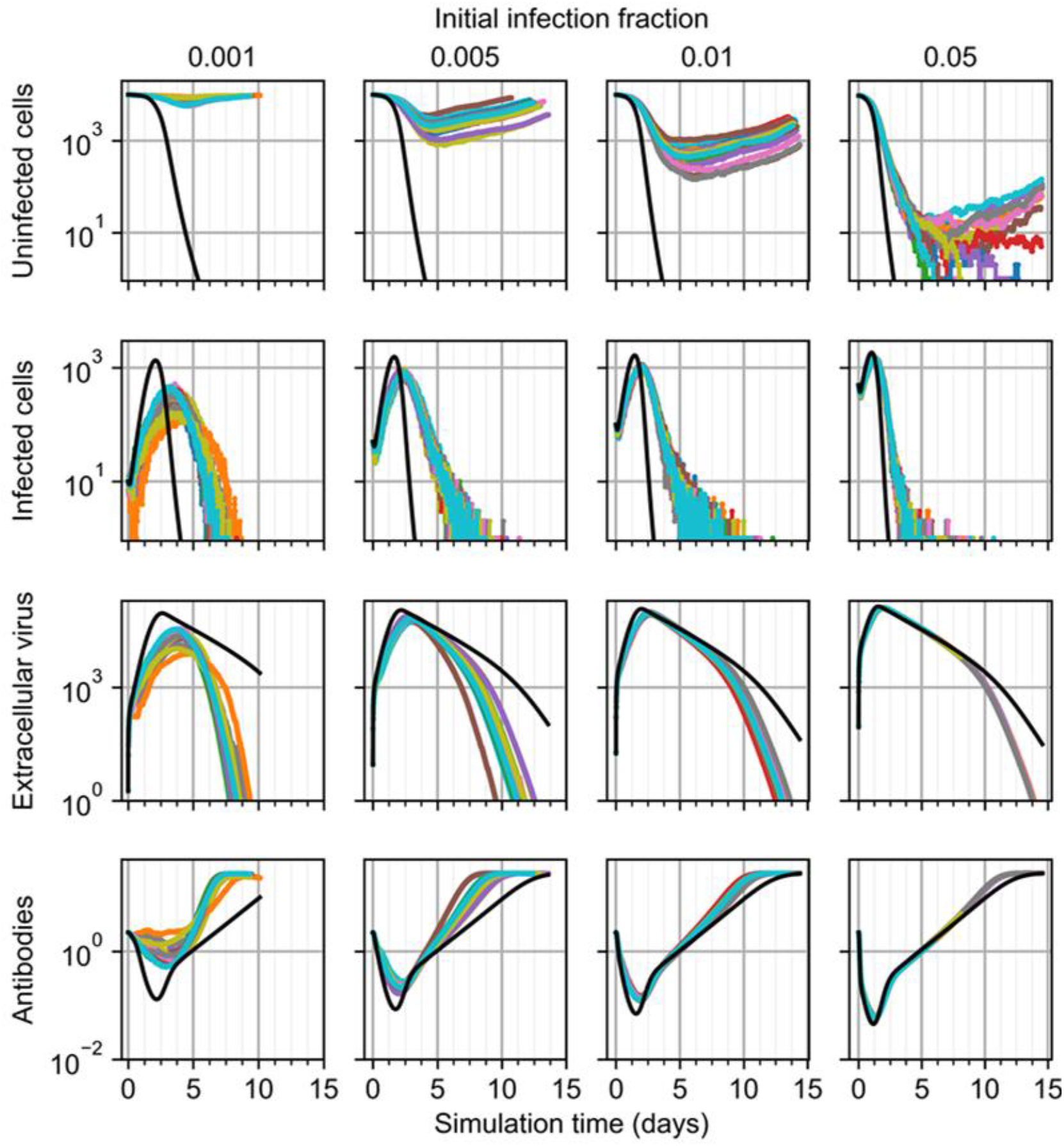
Results from simulation replicas of the spatial model (colored lines) compared to ODE model results (black line) for uninfected cells (top), infected cells (top-middle), extracellular virus (bottom-middle), and antibodies (bottom) for initial infection fraction, from left to right, of 0.001, 0.005, 0.01 and 0.05.

## Discussion

Our simulation scenario (*i*.*e*., the periodic boundary conditions) implies that our simulation replicas are constituent elements of a patterned system, the collection of which the ODE model describes. Comparable work using cellularized models of viral infection have shown that this premise can produce spatial model replicas that are valid spatiotemporal microconfigurations of an ODE model when initially infected cells are essentially spatially homogeneous (22). We showed that simulation replicas using the cellularized model and introduced spatial model mechanisms of this work can also serve as microconfigurations of the original ODE model of influenza infection and immune response under the same boundary and initial infection conditions. However, using spatially homogeneous initial exposure conditions consistently with those employed during calibration of the ODE model generated far less severe infection. This strongly suggests the role of localization of exposure in influenza A infection, in particular that local exposure to virus must be multiple orders of magnitude greater than a uniformly applied exposure to all available susceptible cells. As such, we propose that the particularities of the microenvironment to which the virus is introduced plays a dominant role in disease onset and progression. This is particularly important in therapeutics and modeling, alike, in that spatially resolved patient data of the state of infection may elucidate future health states based on susceptibility of potential lesion sites, which could be better understood and more reliably predicted with spatially-resolved models of the type presented in this work.

Differential adhesion and chemotaxis parameters of the introduced spatial model mechanisms were formulated qualitatively, and only roughly calibrated to recapitulate ODE model results in *Testing Agreement in Small Epithelial Patches*. Interestingly, the employed differential adhesion was necessary to recapitulate ODE model results, the role of which is currently not well-defined. These roles are fairly intuitive when considering the observed temporary aggregation of NK and CD8^+^ T cells in simulations. Were the differential adhesion scheme employed such that NK and CD8^+^ T cells show a preferential attachment to each other, then the observed aggregations at sites of infection due to recruitment would result in ineffective subsequent elimination of infected cells due to the continued aggregation of NK and CD8^+^ T cells. As such, the model predicts that aggregation of local cytotoxic immune cells is due to chemotactic signaling and preferential attachment to infected cells, and that ineffective binding between cytotoxic immune cells makes their subsequent dispersal and translocation, and thus effective contact-mediated local immune response mechanisms, possible.

The most prominent differences between the spatial and ODE models all resolve to localization of type I IFN and recovery. The ODE model, and correspondingly the cellularized spatial model, describe saturated release of type I IFN, the saturation of which is diffusion-limited in the spatial model. This leads to differences not only in total over-production of type I IFN in the spatial models, but also in downstream over-production of virus (*i*.*e*., diffusion-limited anti-viral resistance), with corresponding lesser availability of total antibodies due to interactivity with virus. However, such differences between the ODE and spatial models were shown to be marginal under certain exposure conditions (*e*.*g*., very high initial viral load or infection fraction) and, as previously described, to be significant when the lack of representing localization of virus in the spatial model significantly inhibits the progression of infection (or even its occurrence) in the spatial model epithelial patch. The cellularized Allee effect, which was recast to make associated death and recovery mechanisms dependent on the state and local conditions of individual cells, also produced differences in ODE and spatial model results by allowing recovery with very few total uninfected cells in the spatial model. While we found differences in associated cell deaths to be marginal (and hence, not shown), the spatial model can produce recovery of the epithelial patch in scenarios where associated cell death and a corresponding fatal outcome occur in the ODE model (*e*.*g*., Figure 5), depending on the state and local conditions of uninfected epithelial cells.

### Future Work

The cellularized model of influenza infection and immune response present a number of opportunities for future model development, integration and application. The components of the immune response in the cellularized model can be readily integrated into modular frameworks of viral infection dynamics and immune response that do detailed modeling of other mechanisms like viral internalization and intracellular viral replication dynamics (21). Such activities present two-fold opportunities for novel insights into host-pathogen interactions, in that the immune response components represented here can be leveraged in other viral applications, and likewise integration with other modeling work can be inform further development of the cellularized model presented here. In the case of modeling influenza, detailed modeling of intracellular viral replication while leveraging simulation capabilities like those available in BioNetGen (27) may provide insights into the role of the timing of viral internalization and release, and into the meaning of the calibrated model parameters of the ODE model.

An approach to effectively modeling local features of exposure would significantly improve the ability of our cellularized model to present spatially resolved predictions about the progression of influenza infection and immune response, though will likely require considering mechanisms that are only very implicitly described by the ODE model such as mucociliary clearance. As such, future work should combine the cellularized model presented here with available experimental data and modeling of exposure scenarios. Likewise, future work should further explore and develop a cellular basis for the mechanisms represented by the Allee effect, in particular what all is represented when imposing an organismal-level property like the number of a particular cell type onto the fate of individual cells (*e*.*g*., levels of growth factors, hormones, blood pressure).

The current type I IFN model constitutes the overwhelming majority of computational cost of the spatial model. In particular, calculating a cellular property like resistance *ρ* from the mean value of a local diffusion field requires sufficiently small time steps, since its downstream effects include regulation of future type I IFN production. We plan to make improvements to cellularized mechanisms associated with type I IFN production, as well as to CompuCell3D, to permit larger time steps and better facilitate computational performance. Such improvements are particularly critical to modeling bigger tissue patches and more complicated tissue geometries, and simulating longer scenarios.

Lastly, we are particularly interested in further developing other spatial aspects of the cellularized model to further elucidate associated cellular and spatial aspects of influenza infection and immune response, as well as the immune response, in general. We can use *in vitro* data of cell migration to refine the proposed locomotion model of local immune cells at a site of infection, both to better understand how adhesion and chemotaxis affect the effectiveness of the immune response, and to isolate necessary additional model mechanisms to better represent local aspects of immune response. At a broader level, we can perform similar activities to this project but for other sites of interest associated with the ODE model. For example, the ODE model presents systemic response data that can guide development of spatiotemporal, cell-based models of B cell maturation, antigen presentation, and antibody production and circulation. We envision a computational framework consisting of multiple compartments simulating spatiotemporal models of various sites of interest throughout an organism (*e*.*g*., multiple sites of infection, lymph nodes, thymus), which could be interconnected using the techniques of cellularization in similar fashion to what was employed in this work.

## Conclusion

In this work we developed and employed a multiscale, spatiotemporal, stochastic, cell-based model of influenza infection and immune response by cellularization of an existing, calibrated ODE model. We developed spatial models of necessary mechanisms related to differential adhesion and chemotaxis of local immune cells to recapitulate ODE model results using the spatial model, and generated a cellularized form of the Allee effect to describe recovery of epithelial cells in terms of cell state and local conditions. We used the developed spatial model to show how exposure to virus should be locally concentrated to generate significant infection in an epithelium, while uniform exposure to virus is likely ineffective. We also used our developed spatial models to elucidate the necessary roles of differential adhesion and local chemotaxis for an effective local immune response, both concerning the locating by macrophages, and the eliminating by NK and CD8^+^ T cells, of infected cells.

## Acknowledgments

TJS and JAG acknowledge funding from National Institutes of Health grants U24 EB028887 and R01 GM122424 and National Science Foundation grant NSF 1720625. GBE acknowledges funding from National Science Foundation grant NSF 1951099. This research was supported in part by Lilly Endowment, Inc., through its support for the Indiana University Pervasive Technology Institute. The funders had no role in manuscript preparation or the decision to submit the work for publication.

## Supplementary Materials

## Appendix 1

This section presents the complete cellularization of the ODE model. The cellularized form of each ODE is presented along with the original ODE form according to the cellularization scheme described in *Models and Methods*. In general, we employ the notation that for a number of cells *Z* of cell type 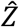 according to the scale of the ODE model, the corresponding number of cells of type 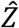 in the spatial domain is written as Z. The same notation is employed for soluble signals (*e*.*g*., *A* measured at the scale of the ODE model is *a* measured at the scale of the spatial model), with the exception that we write *T*^∗^ as the total amount of TNF in the spatial domain (so as not to be confused with time). For the soluble signals chemokines *C*, IL-10 *L*, type I IFN *F* and extracellular virus *V* with explicitly represented spatial distributions 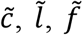 and 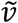, respectively, the global amounts *c, l, f* and *v* are calculated as the volume integral of their respective spatial distribution over the spatial domain. For all other soluble signals, their spatial distribution is treated as uniform over the spatial domain when appropriate (*e*.*g*., for antibodies, *A* = *η*^−1^ *a*= *θ* ^−1^ *ã*).

The ODE model describes two pro-inflammatory signals Σ_1_ and Σ_2_, the first of which describes dead cells and TNF as stimuli for production of chemokines and IL-10 and stimuli to the second signal, and the second of which describes the first and extracellular virus as stimuli for production of TNF. For the first signal,

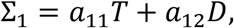

and for the second signal,

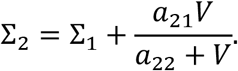

Both signals are treated as global signals in the spatial model with corresponding values *σ*_1_ = ηΣ_1_ and *σ*_2_ = ηΣ_2_, such that for the first signal,

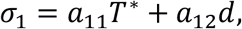

and for the second signal,

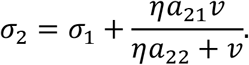

### Soluble Signals

The model mechanisms for chemokines, IL-10, type I IFN and extracellular virus are described in *Models and Methods*, where also a cell-based description of viral resistance *ρ* due to type I IFN is described. In the ODE model, the corresponding model of viral resistance *R* of the epithelial cell population takes the form,

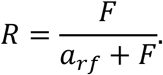

The rate equation for extracellular virus has the form,

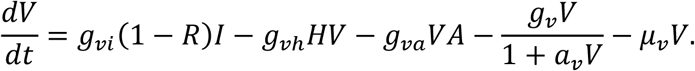

The rate equation for chemokines in the ODE model has the form,

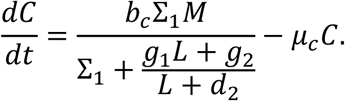

The rate equation for IL-10 in the ODE model has the form,

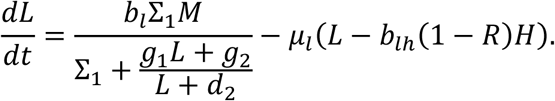

The rate equation for type I IFN has the form,

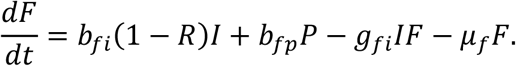

In the inflammatory response, the ODE model describes production of TNF by the second pro-inflammatory Σ_2_ and macrophages, and regulated by IL-10,

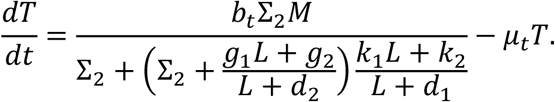

The spatial model treats TNF as uniformly distributed, with the corresponding rate equation,

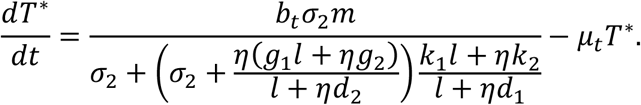

The ODE model describes production of ROS by tissue neutrophils and its uptake by uninfected and infected cells,

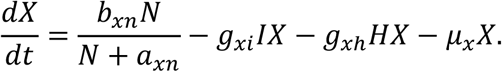

The spatial model treats ROS as uniformly distributed, with the corresponding rate equation,

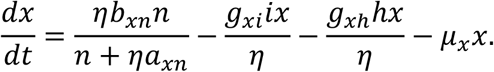

In the immune response, the ODE model describes production of IL-12 by APCs mediated by CD4+ T cells,

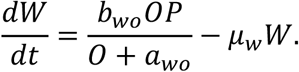

The spatial model treats IL-12 as uniformly distributed, with the corresponding rate equation,

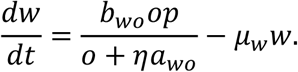

The ODE model describes production of type II IFN by NK and CD4+ T cells mediated by IL-12,

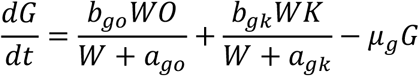

The spatial model treats type II IFN as uniformly distributed, with the corresponding rate equation,

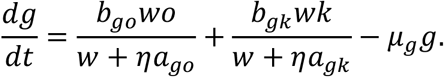

The ODE model describes production of antibodies by B cells, their reaction with extracellular virus, and a homeostatic nonzero level,

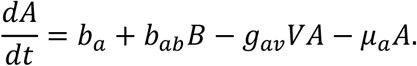

The spatial model treats antibodies as uniformly distributed, with the corresponding rate equation,

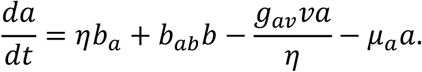

### Epithelial Cell Dynamics

In the epithelial sheet, the ODE model describes, in order, an Allee effect, infection and killing by ROS, in the dynamics of the uninfected cell population,

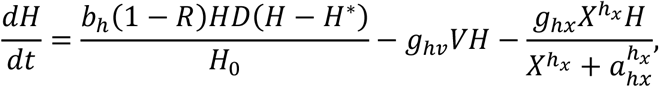

where *H*^∗^ is a critical population value above which recovery occurs, and below which additional death of uninfected cells occurs. In the dynamics of the infected cell population, the ODE model describes, in order, infection and killing by ROS, NK cells, CD8+ T cells and resistance-associated effects,

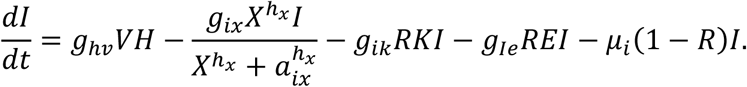

The Allee effect is split into the discrete, stochastic processes of uninfected cells becoming dead cells, and of dead cells becoming uninfected cells. Both processes occur due to neighborhood conditions of individual cells. Using the notation 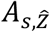 to denote the contact area between cell *s* and 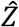 − type cells and *A*_*s*_ to denote the total contact area of cell *s*, the following equalities are true under well-mixed conditions,

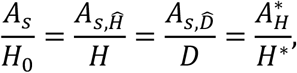

where 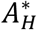 is a critical contact area with uninfected cells above which dead cells can recover, and below which uninfected cells can die. The probability of recovery in each dead cell *s* then occurs according to an equipollent recovery rate *a*_*H*_ = *a*_*H*_(*s*),

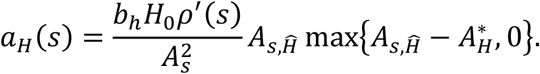

Likewise the probability of death of an uninfected cell due to the Allee effect occurs according to an equipollent Allee death rate *a*_*D*_,

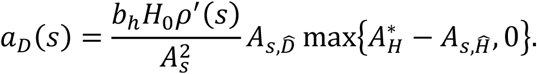

Noting that for epithelial cells, 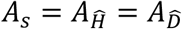, and using the contact-mediated equipollent rate *γ* from cellularization, the recovery and Allee death rates can then be written as

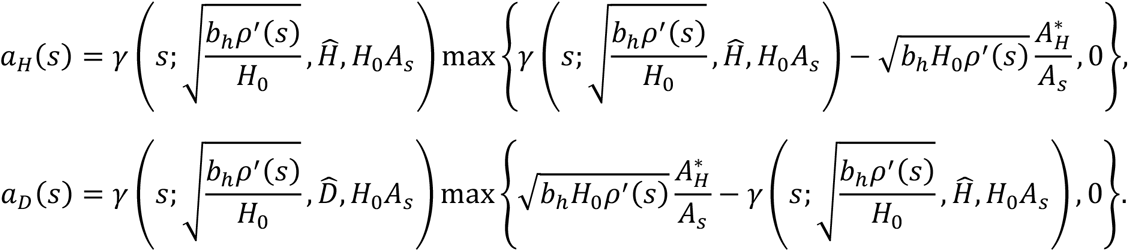

Applying these cellularized forms of the Allee effect and the other mechanisms described in the ODE model, the probability of infection and death for each uninfected cell are, respectively,

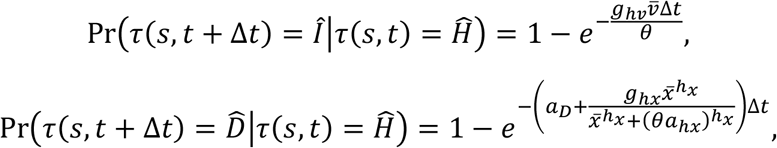

where *g*_*hv*_, *g*_*hx*_ and *a*_*hx*_ are ODE model parameters. The probability of death for each infected cell is then

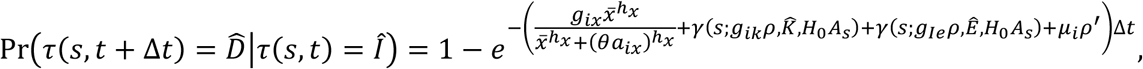

where *g*_*ix*_, *a*_*ix*_, *g*_*ik*_, *g*_*Ie*_ and μ_*i*_ are ODE model parameters. The probability of recovery for each dead cell is then

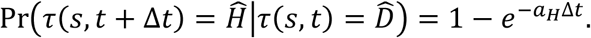

### Immune Cell Dynamics

In the inflammatory response, the ODE model describes recruitment of macrophages by chemokines and a resident macrophage population,

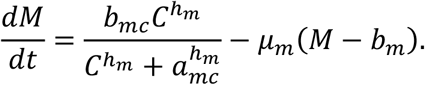

The spatial model explicitly models macrophages in the spatial domain. Inflow of macrophages is described by the cellularized probability,

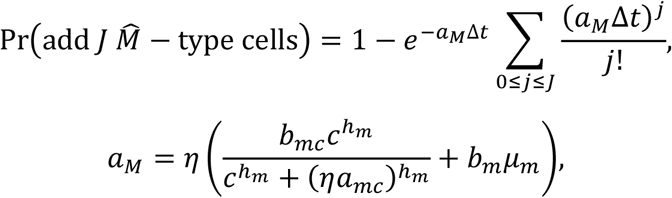

and outflow of macrophages is described by the cellularized probability,

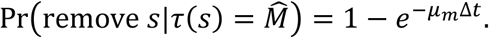

The ODE model describes recruitment of blood neutrophils by TNF regulated by IL-10 and recruitment of tissue neutrophils from blood neutrophils by chemokines. For blood neutrophils according to the ODE model,

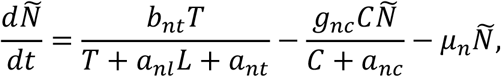

and for tissue neutrophils according to the ODE model,

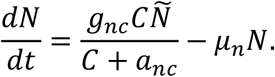

The spatial model treats both blood and tissue neutrophils as globally acting populations. As such, in the spatial model blood and tissue neutrophils are also modeled with an ODE scaled to the size of the spatial domain. In the spatial model, for blood neutrophils,

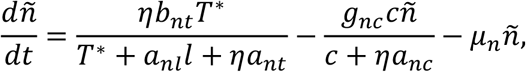

and for tissue neutrophils,

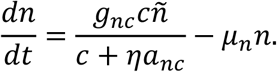

In the immune response, the ODE model describes the generation of APCs by extracellular virus and dead infected cells *D*_*I*_ upregulated by type II IFN and a resident population,

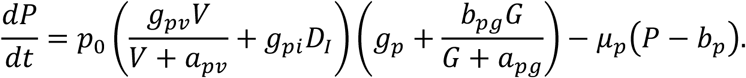

The spatial model describes APCs as globally acting with a scaled ODE model,

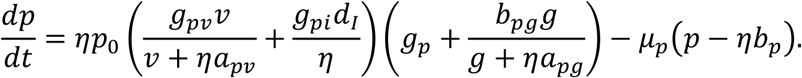

The ODE model describes the recruitment of NK cells by chemokines regulated by resistant infected cells and a resident population,

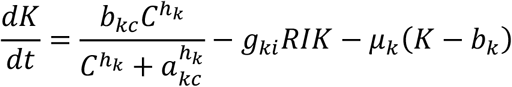

The spatial model explicitly models NK cells in the spatial domain. Inflow of NK cells is described by the cellularized probability,

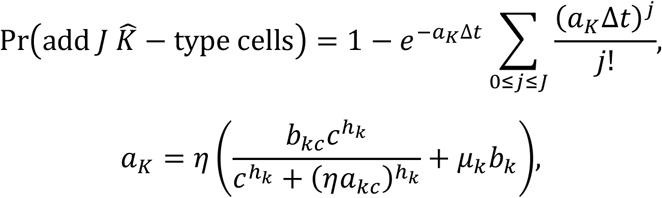

and outflow of NK cells is described by the cellularized probability,

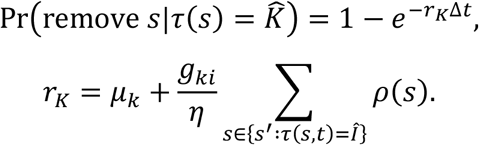

Note that the effects of infected cells on the NK cell population could also be interpreted as resulting from NK cell death by contact-mediated interactions, as with killing of infected cells by NK cells. We instead implement the simpler case here and assume that the infected cell population has an indirect, inhibitory effect on recruitment of NK cells to the local domain. The ODE model describes the recruitment of CD4+ T cells by APCs,

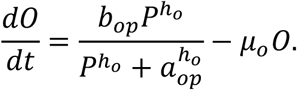

The spatial model describes CD4+ T cells as globally acting with a scaled ODE model,

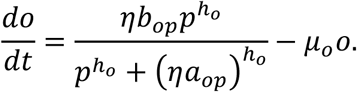

The ODE model describes the recruitment of CD8+ T cells by APCs regulated by resistant infected cells,

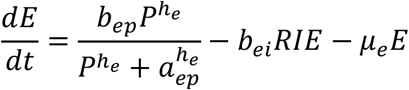

The spatial model explicitly models CD8+ T cells in the spatial domain. Inflow of CD8+ T cells is described by the cellularized probability,

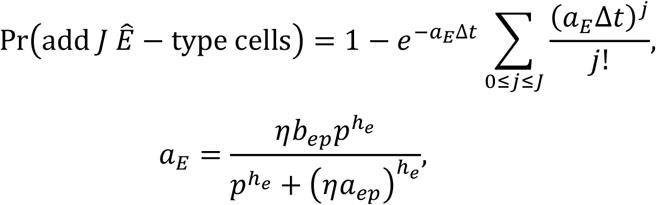

and outflow of CD8 + T cells is described by the cellularized probability,

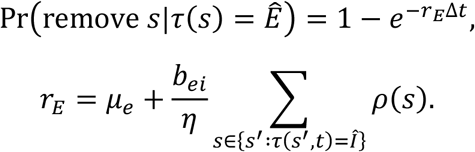

As with NK cells, that the effects of infected cells on the CD8+ T cell population could also be interpreted as resulting from CD8+ T cell death by contact-mediated interactions. We also implement the simpler case for CD8+ T cells and assume that the infected cell population has an indirect, inhibitory effect on recruitment of CD8+ T cells to the local domain. The ODE model describes the recruitment of B cells by the combined action of IL-12 and APCs and a resident population,

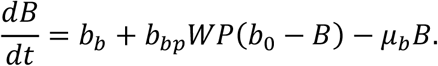

The spatial model describes B cells as globally acting with a scaled ODE model,

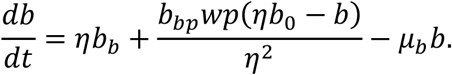

## Appendix 2

**Table A2.1.**
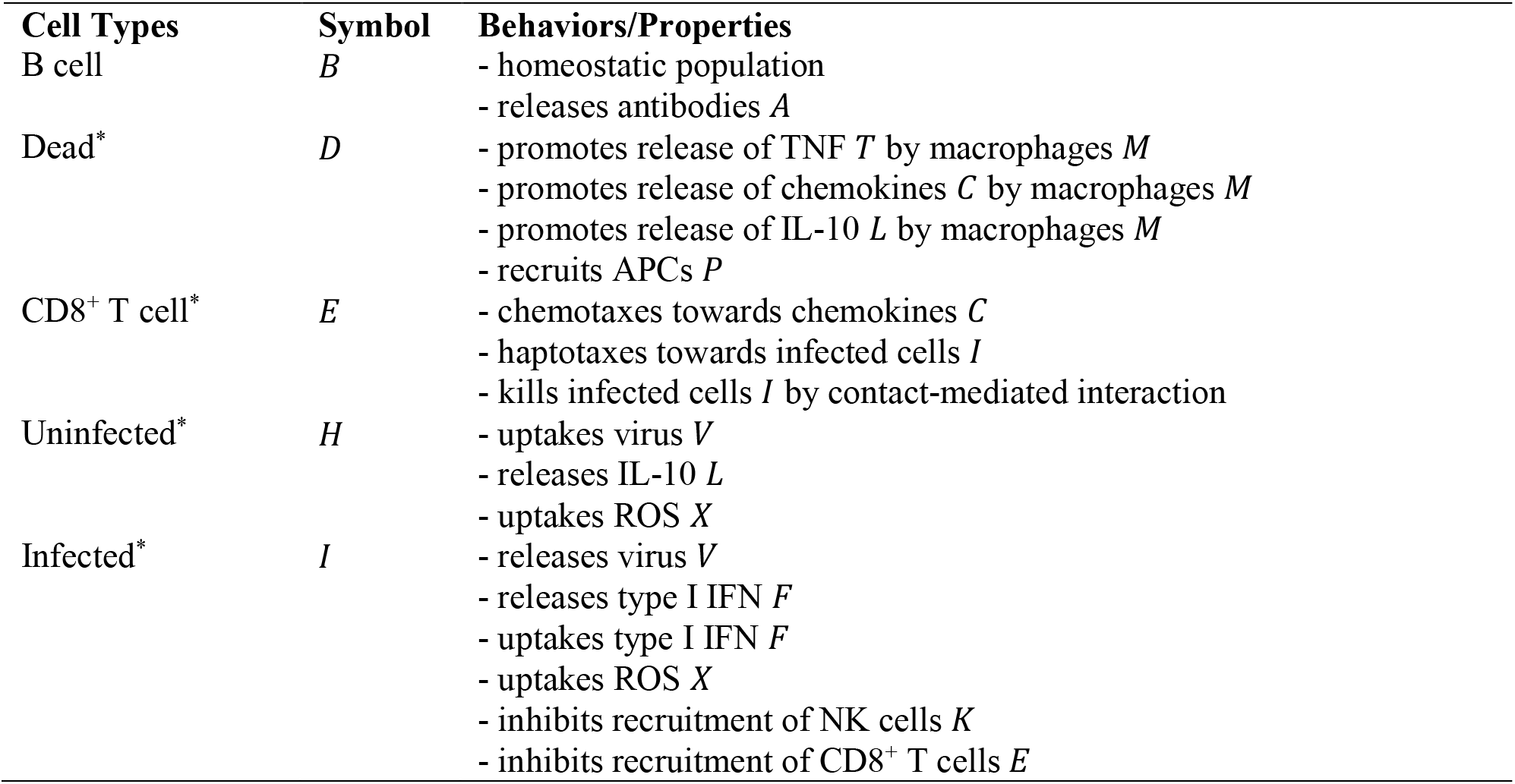
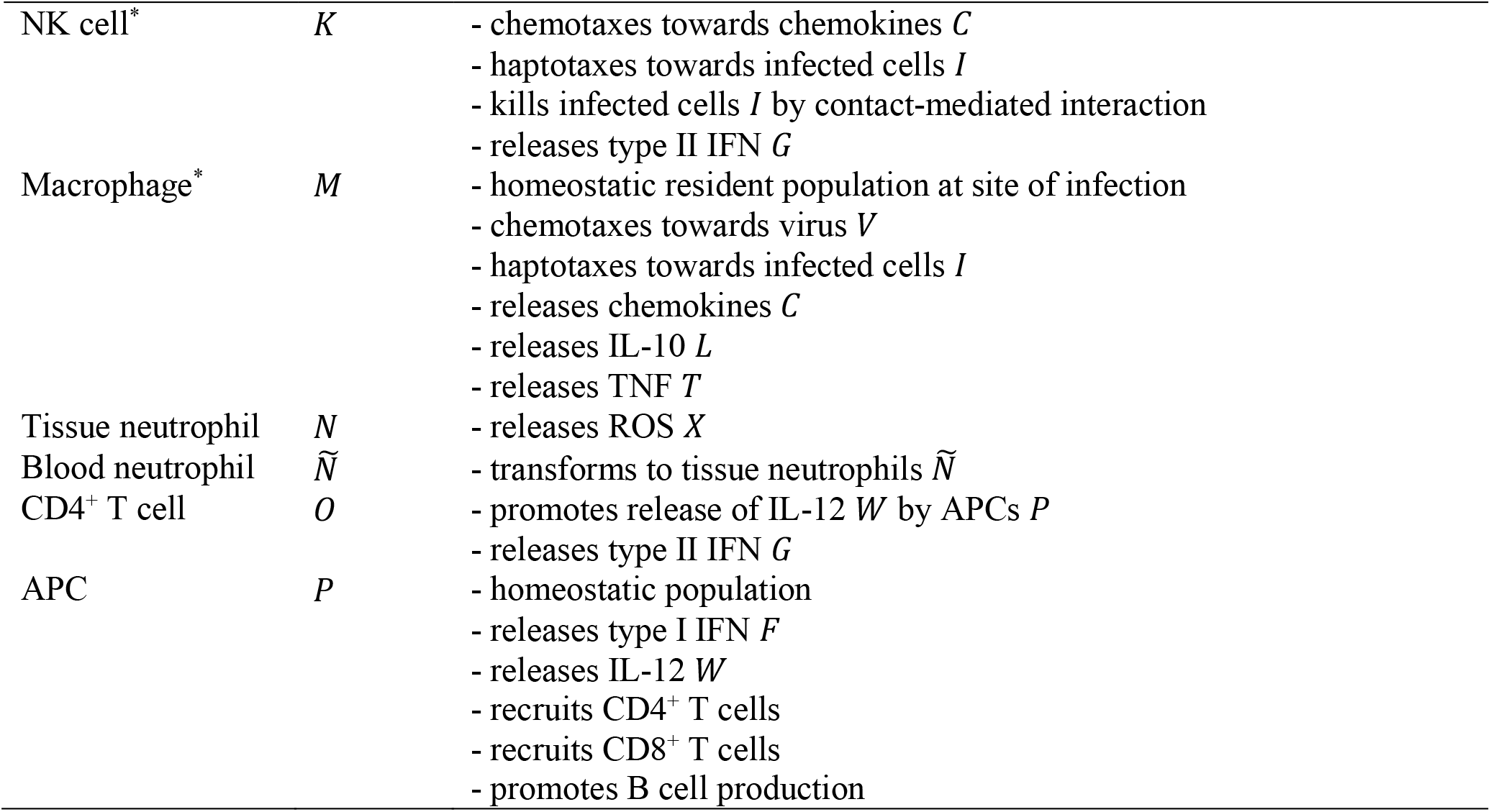
Cell types and their mathematical symbols, behaviors and properties in the model. Locally modeled cells are denoted with (^*^).

**Table A2.2.**
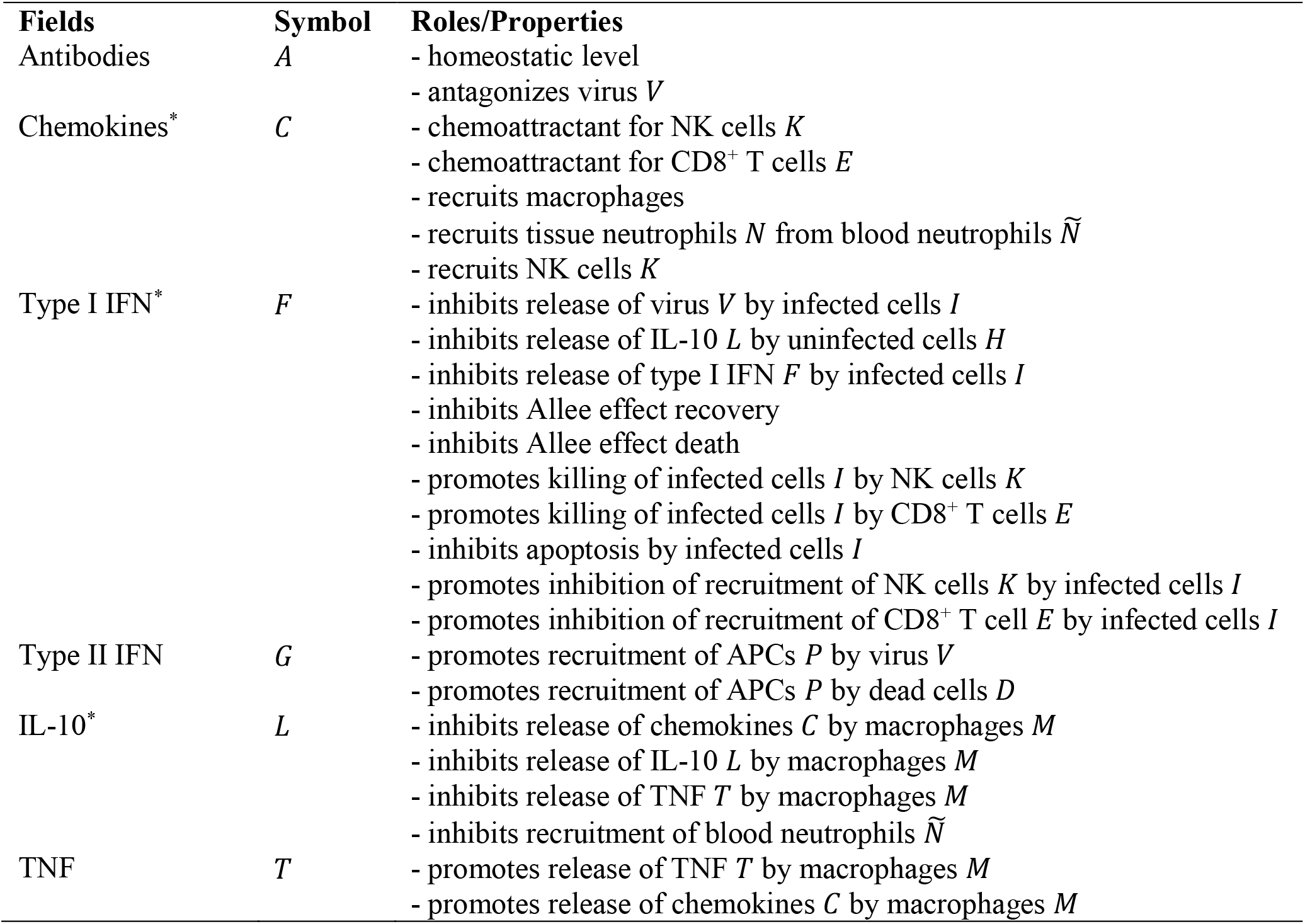
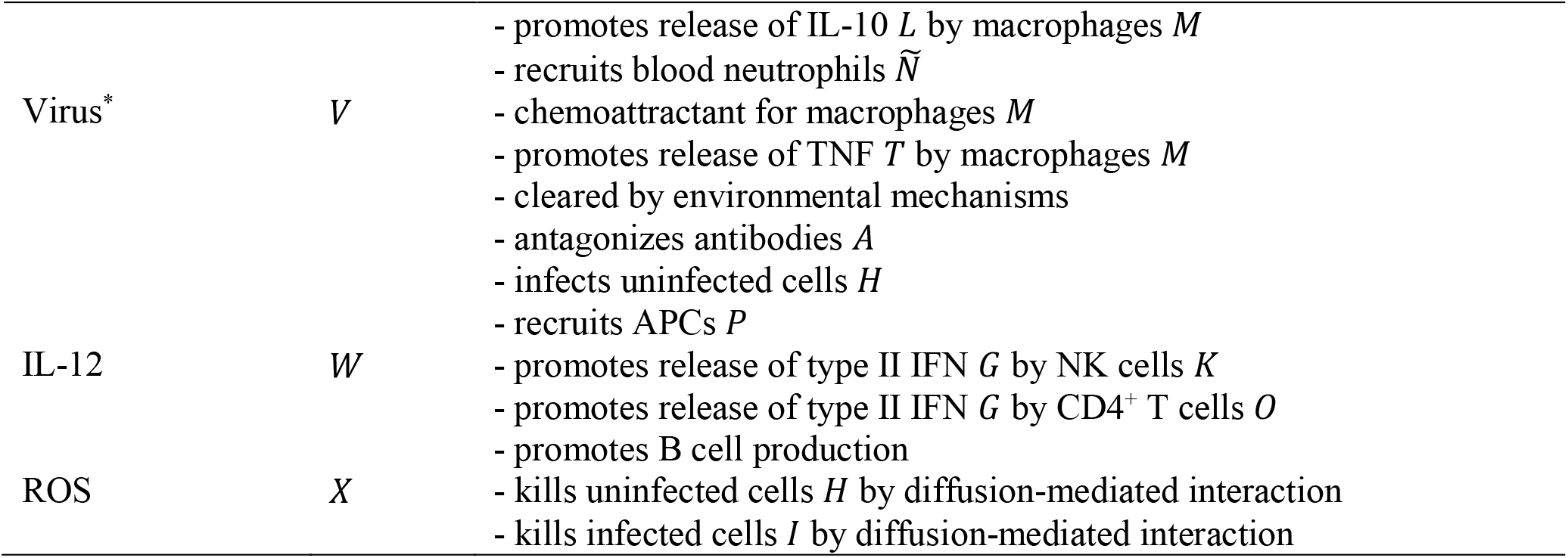
Fields and their mathematical symbols, roles and properties in the model. Locally modeled fields are denoted with (^*^).

